# High Intensity Interval Training in Aged Female Mice Preserves Physical, Cognitive, and Cardiovascular Function

**DOI:** 10.64898/2026.06.07.730494

**Authors:** Drew Theobald, Phillip Williamson, Alexandra Johnston, Lucas Tripp, Ayodeji A. Olabiyi, Xavier Silvers, Ashby Dickerson, Tuan D. Tran, Lisandra de Castro Braz, Srinivas Sriramula, Ted G. Graber

**Author notes:** Corresponding/Senior Author: Ted G. Graber, PhD, Assistant Professor East Carolina University, Dept. of Physical Therapy. Author Emails: Drew Theobald, MS, Phillip Williamson, BS, Alexandra Johnston, MS, Lucas Tripp, DPT, Ayodeji A. Olabiyi, PhD, Xavier Silvers, BS, Ashby Dickerson, BS, Tuan D. Tran, PhD, Lisandra de Castro Braz, PhD, Srinivas Sriramula, PhD, DVM.

## Abstract

**BACKGROUND:** Along with advancing age comes declines in physical, cognitive, and cardiovascular function. This diminished capacity may lead to decreased ability to perform activities of daily living, disability onset, and loss of independence. Exercise is a regenerative medicine therapy that can mitigate this loss of function. High intensity interval training (HIIT) is an aerobic exercise paradigm consisting of intense activity periods interspersed with bouts of active recovery. Previously we demonstrated that HIIT preserved physical function in adult, middle-aged, and older male mice. However, whether HIIT preserves physical, cognitive, and cardiovascular function, mitigates frailty, and improves brain and heart health in older adult female mice remains unknown.

**HYPOTHESIS:** Cognitive, physical, and cardiovascular function in older adult female C57BL/6 will be preserved in exercised mice (HIIT) versus sedentary control (SED).

**METHODS:** Mice (HIIT and SED, both n=9, 24m at end) were tested pre/post-intervention for physical (rotarod, treadmill, grip meter, inverted cling, voluntary wheel running, activity monitor), cognitive (open field, novel object recognition, puzzle box, y-maze), and cardiovascular (blood pressure, echocardiogram) function, body composition, and whole body calorimetry. The mice underwent 14-weeks of HIIT training with progressive volume and intensity.

**RESULTS:** HIIT significantly (p<0.05) increased or preserved function in many tests including: aerobic capacity (+71% HIIT versus, vs, no change, NC, in SED), four limb strength/endurance (−67% SED vs -28% HIIT), forelimb strength (−16% SED vs NC HIIT), overall motor function (NC SED vs +39% HIIT), executive function (NC SED vs +73% HIIT), and exploratory behavior, which improved across multiple tests with HIIT while remaining unchanged in SED. HIIT also reduced both systolic blood pressure by 12% (−17 mmHg) and mean arterial pressure by -16 mmHg. In addition, HIIT significantly reduced cardiac fibrosis, increased muscle fiber type 2a percentage, reduced IL-1β expression in the hypothalamus, and mitigated frailty onset.

**CONCLUSION:** HIIT significantly reduced age-related functional loss in all three domains assessed while preventing frailty onset in older adult females and improving markers of brain and heart health.

## Introduction

With aging comes the onset of chronic diseases and loss of functional capacity. The primary risk factor for many chronic diseases is advancing age, with 90% of older adults having at least 1 chronic illness, and nearly half having multi-comorbidities.(1,2) The global population of older adults is advancing at an exponential rate, with projections of an increase from roughly 700 million adults over the age of 65 in 2019 to more than 1.5 billion by 2050.(3) Healthspan is the period of life with a relatively low burden of disease and adequate functional capacity to perform activities of daily living and remain independent. Extending healthspan, not necessarily lifespan, is a critical concern as the population of older adults continues to grow. Thus, interventions that slow the aging process, increase resilience, preserve intrinsic capacity, and maintain physiological function as long as possible are critical for helping society sustain essential safety net programs, such as social security and Medicare in the United States.(4)

Age-related functional decline to some degree is an inevitable consequence of getting older; however, the rate and severity of that decline is modifiable—potentially though lifestyle changes such as by increasing activity and eating a healthy diet. Geroscience as a discipline seeks to use medical advances to slow the aging process, and thus retard the downstream effects of age-related diseases, syndromes, and conditions including sarcopenia (age-related loss of muscle mass and strength), frailty (inability of body to maintain homeostasis and respond to challenges), and cognitive decline or dementia. Exercise in its varied forms is a therapy to preserve functional aptitude; however, the exact molecular basis of the beneficial effects is not fully understood within the context of aging physiology, nor is the optimal volume, intensity and/or types of exercise needed to maximize benefits. Developing preclinical models to understanding mechanisms by which exercise preserves function could lead to novel therapeutic targets and exercise mimetics.

High intensity interval training (HIIT) is an exercise paradigm consisting of periods of intense activity interspersed with periods of active recovery or resting. HIIT has been demonstrated to preserve physical function and mitigate frailty in both female and male mice.(5–7) Endurance training, including HIIT, can improve measures of cardiovascular health, including systolic hypertension.(8) Physical, cardiovascular, and cognitive function, as well as muscle, heart, and brain health, are interlinked, and we believe that an intervention designed to improve one will also improve the others. Mechanisms through which exercise may influence multiple domains include circulating factors, such as exerkines and extracellular vesicles, emitted by multiple organs and tissues, not just skeletal muscle, during and after an exercise session.(9–11) Furthermore, the hallmarks of aging include elevated global inflammation, i.e. “inflammaging”, loss of proteostasis, and mitochondrial dysfunction, among others.(12) Exercise is a potential mediator of these hallmarks, and may act as a potent geroscientific anti-aging intervention.(13)

Previously, our group showed that *low volume* HIIT preserved physical function in both adult and older adult male C57BL/6 mice, but was less successful in promoting improvement of overall physical or cognitive ability in early middle-aged males, albeit aerobic capacity was greatly increased compared to a sedentary group.(5,14) However, whether HIIT preserves physical, cognitive, and cardiovascular function, as well as brain and heart health, in older adult female mice remains unknown. Previously our lab demonstrated a markedly different functional aging trajectory between males and females, diverging particularly beginning in middle-age.(15) Coupled with these different patterns of functional loss with aging, and since we also know that there are divergent exercise effects based on sexual dimorphisms, it is critical to ensure studies involve females.(15,16)

In this current study, we set out to determine the effect of HIIT on the physical, cognitive, and cardiovascular function of older adult female mice. We further sought to investigate potential effects on heart, muscle, and brain health in the context of fibrosis, inflammation, and myocyte morphology. These studies will provide insight into whether HIIT promotes healthy aging in older females by preserving functional capacity and improving cardiac, skeletal muscle, and brain health. To our knowledge this is the first such comprehensive study of HIIT in older female mice.

## Methodology

### Animals

We obtained female C57BL/6J mice from the Jackson Laboratory, n=24 at age 12m (months). An additional young adult control group (3 months of age, n=4) was used for some of the immunohistochemical experiments. Mice were group-housed under 12-h light/ 12-hr dark cycles at 22 °C, with *ad libitum* access to food and water. The mice were treated humanely in accordance with East Carolina University Institutional Animal Care and Use Committee (IACUC) guidelines and our approved animal use protocol.

### Study Design

Mice (n=24) were aged to the start of the training study at 18 months and underwent pretesting for function. We randomly assigned mice to either the sedentary control group (SED) or the exercise training group (HIIT). Mice trained for 14 weeks and then euthanized at 24 months for tissue collection following post-testing for function. See **Figure 1 for the study design.**

**Figure 1.**
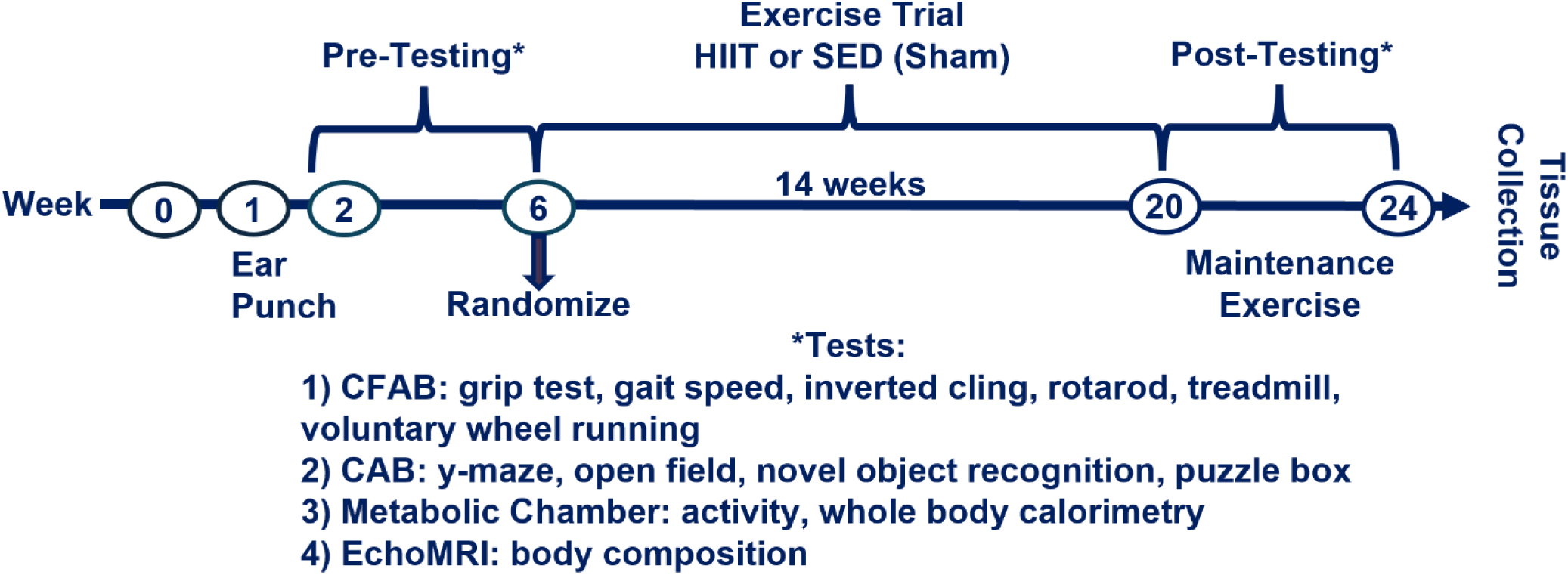
Study Design. KEY: HIIT=mice undergoing 14 weeks of high intensity interval training, SED = sedentary control group with a sham treatment. CFAB = comprehensive functional assessment battery, CAB = cognitive assessment battery. Maintenance exercise was performed to maintain adaptations during the post-testing period. Tissue was collected and mice euthanized two days after their final maintenance exercise session.

### Training

Our typical individually personalized training for HIIT has been previously described.(5,14) In brief, with differences from previous studies as follows: During pretesting for function, mice underwent the treadmill maximum speed (max speed) test for aerobic capacity. The baseline to set the HIIT intervals was the maximum obtained speed (meters/minute, m/min). The first week of training was acclimation where the mice ran for three intervals (one at 70%, the next at 75%, and the last again at 70% of max speed). Each training session began with a 2-minute walk at 5 m/min, then the first high intensity interval began with a 30 second accelerating ramp-up to 70% of max speed, then maintained this running speed for 60 seconds, followed by a 30-second ramp-down deceleration to walking speed (5 m/min). Then the mice completed 60 seconds of active recovery at walking speed prior to the start of next interval. After completion of all intervals, the mice had a 2-minute cool down at walking speed. Both volume (increased number of intervals) and intensity (increased running speed) periodically increased over the 14-week training period. At the ½ way point (7 weeks) the mice reperformed the max speed test and the interval speeds then adjusted if needed. At the end of the training study mice ran for 7 intervals of 70-75-80-80-80-80-75% of max speed (from the mid-point measure). During the functional post-testing, the mice ran maintenance training 2-3 times per week, at similar intensity and volume to the ending training (described in the previous sentence), to maintain fitness adaptations.

The SED mice did a sham control in which they were placed on the treadmill for the same duration as the HIIT group, but the treadmill ran for only 5 seconds at 5 m/min once every 3 minutes. This sham protocol provided similar enrichment and stimulation to HIIT, minus exercise stimulation, which was a critical component to match for the purposes of cognitive function testing.

### In vivo Outcome Measures

#### Physical Function

We previously described and validated our composite scoring system, CFAB (Comprehensive Functional Assessment Battery) in male and female mice of various ages in both cross-sectional and longitudinal studies.(5,14,17–19) CFAB consists of five well-validated tests for multiple domains of physical function and exercise capacity including rotarod (overall motor function including elements of endurance, balance, coordination, and power production), inverted cling (four-limb strength/endurance), grip meter (fore-limb strength), treadmill max speed test (endurance and aerobic capacity), and voluntary wheel running (VWR, volitional exercise rate), with details on running the tests reported previously.(19) Briefly, in this study, we used pre- and post-CFAB scores, as well as the change in CFAB scores (ΔCFAB), to compare exercise efficacy and changes in function over time. We compared each of the five determinants for every mouse (pre- and post-intervention) to the mean score for the overall pre-intervention cohort and assigned a value equal to the number of standard deviations away from the cohort mean. For example, a rotarod score of x was compared to the overall mean of y and, whether positive or negative, the number of standard deviations (of the overall pre-test for all mice) from y became that score for the rotarod. We treated the other determinants the same way and then summed them all to comprise the CFAB composite score-with a more negative score indicating lower function.

In addition, we also measured x/y/z plane activity using lasers and voluntary wheel running in the Promethion metabolic cages. We took the total activity (all meters moved in x/y/z planes) and wheel measures (total meters, m, run), divided by the number of hours the mice were in the cages and then multiplied by 24 to get mean per day.

*Cognitive Function,* CAB (Cognitive Assessment Battery): For cognitive function we used our CAB determinants of puzzle box (executive function and memory), y-maze (working memory and exploratory behavior), open field (exploratory behavior and anxiety), and novel object recognition (NOR, long term memory and exploratory behavior). Details about the tests have been previously detailed, but with this older group of mice we made some changes.(14) The amount of time allowed for each of the tests was increased since we noticed that the older mice had slower starts (ten minutes for open field and NOR). Since a small subset mice do not complete some challenges during the puzzle box, we assigned a number to a mouse that was unable to complete a task as being the maximum time allotted for the test (example, 6 minutes) plus 30 seconds (score if task not completed = 6:30). Lighting was lowered to approximately 40-50 lux (measured at all 4 corners in the open field box, or all 3 arms in the y-maze), to better encourage exploratory behavior, except for the puzzle box challenges where the light was set at ∼500 lux (where a bright open field box is incentive to enter the escape tunnel and dark escape box). We utilized AnyMaze software to analyze the data recorded on a GoPro12 camera. See Online **Supplementary Methods** for further details.

We converted CAB into a composite score by determining which quartile an individual mouse was in for each of our included measures: open field total distance and center time, NOR discrimination ratio and total exploration time, puzzle box total time (challenge 1 and 2) and total time challenge 3, and y-maze total alternations and spontaneous alternation ratio, then summed these to get a total score, with higher score equal better performance. Note that in the puzzle box tests quartiles were in reverse order since lower numbers were better (i.e., quartile 1 was the best, so it was converted to 4 to match the other tests), and in the other three tests higher numbers were better (i.e., quartile 4 is best).

#### Frailty

We measured frailty based upon our prior work.(20,21) Note that CFAB correlates very highly with Frailty Score and we can use CFAB scores as a proxy for frailty scores.(17,22) Briefly, mice with a determinant score (e.g., treadmill time in seconds) below the cutoff point (≤ 1.5 standard deviations (sd) below the baseline mean of all mice pre-intervention) of the given determinant received 1 point, zero otherwise. We did this for each determinant (VWR, inverted cling, grip strength, treadmill max speed test, and rotarod) and then summed the points earned. Any mice with 3 or more points equaled frail, two considered pre-frail, and less than 2 as non-frail. We compared pre- and post-training Frailty scores (post-training frailty used the pre-intervention cut-off) and compared CFAB to Frailty. Additionally, we compared the post-tests using a modified version of the mouse frailty index, as a 6^th^ determinant as we recently published.(23–25) Further details on the frailty measurement are in the online **Supplemental Methods** section.

#### Blood pressure measurement

We used a noninvasive tail-cuff plethysmography according to the manufacturer’s protocol (CODA, Kent Scientific), described in further detail in the Supplemental Methods section. Mice acclimatized to the procedure before baseline measurements and then blood pressure readings taken for three consecutive days each week. We report the mean values, representing one weekly measurement for each mouse.

#### Echocardiographic Imaging

We assessed systolic and diastolic function before and after the intervention in both the HIIT and SED groups with echocardiographs, using a digital ultrasonic imaging system (Vevo 3100, Fujifilm VisualSonics, USA), as described previously and according to the Guidelines for Measuring Cardiac Physiology in Mice.(24–26) Further details are in the **Supplemental Methods**.

#### Body composition

We measured body mass weekly during training, and body composition by EchoMRI before and after the intervention. Lean mass and fat percentage changes reported as described previously.(5,15)

#### Whole-body metabolism

We used Promethion metabolic chambers (Sable Systems) according to the manufacturer’s instructions, to measure and analyze voluntary wheel running, x/y/z-plane activity, oxygen consumption, carbon dioxide production, respiratory exchange ratio, and energy expenditure (see **Supplementary Methods** section for further details).

#### Immunohistochemistry

We processed brain, heart, and soleus muscle tissues for immunofluorescence staining, acquiring images using fluorescence microscopy, and quantifying mean fluorescence intensity with ImageJ. See **Supplementary Methods** Section for further details. Briefly:

#### Glial cell morphology analysis

We assessed microglial morphology in IBA1-labeled brain sections using confocal microscopy and Fiji/ImageJ skeleton analysis, as described previously.(27) Astrocyte complexity quantified using Sholl-based ramification analysis in Fiji/ImageJ.

#### Cardiomyocyte cross-sectional area

For cardiomyocyte cross-sectional area analysis, we labeled cell membranes with wheat germ agglutinin (WGA) conjugated to Alexa Fluor 488 (Invitrogen W11261; 1 mg/mL stock, 1:200 dilution) for 10 min as previously described.(26,27) Sections were mounted with VECTASHIELD antifade mounting medium with DAPI. We acquired images using a fluorescence microscope (Keyence/Echo Revolve), and cardiomyocyte cross-sectional area quantified in ImageJ by a blinded investigator.

#### Measurement of collagen

To determine the extent of collagen deposition as a measure of cardiac remodeling, we stained heart tissue sections with 0.1% Picrosirius Red to visualize fibrillar collagen as previously described.(28) Sections were then imaged under standard brightfield microscopy by an investigator blinded to the experimental groups, and collagen-positive staining quantified using identical thresholding parameters across all groups.

#### Soleus muscle morphology

We assessed soleus muscle fiber type distribution and cross-sectional area by immunofluorescence in a randomly selected subset of mice, as described previously.(30,31) Specific details and antibodies used are in the **Supplemental Methods** section.

#### Statistical analysis

Data reported as means ± standard error (SE), standard deviation (SD), effect size (Hedges’ g and Cohen’s d, or η2), with statistical significance set at p<0.05, and trends when 0.05≤p≤0.10. We considered a strong effect size as Hedge’s g > 0.70 (Hedge’s correction to Cohen’s d used due to the small n). We report any potential issues with normality by reporting skew, kurtosis, and the results of the Kolmogorov-Smirnov and Shapiro-Wilks tests in the **Supplemental Data Sets** for each of the functional variables. We then ran non-parametric tests (Wilcoxon Ranked Sum for paired data and Mann-Whitney U-test for independent data and/or Independent Samples Median Test) if we determined a need based upon the above criteria. We used SPSS v31 (IBM), AnyMaze, and GraphPad Prism (10.4.0 GraphPad Software) for statistical analysis. Pre- and post-measurement means with 2 groups were compared within groups by the paired t-test, and between groups compared using the independent samples t-test. Means from three groups were compared with mixed-model ANOVA (for example, treadmill speed: 2x3 ANOVA consisting of 2 groups, SED and HIIT, with three timepoints, pre-, mid-, and post-intervention).

Immunofluorescence statistical analyses used GraphPad Prism (v. 10.4.0, GraphPad Software) for brain and heart data, and SPSS v31 for muscle data. All experiments and analyses were performed in a blinded fashion by two independent investigators. For comparison of HIIT, SED, and 3m control, we utilized one-way ANOVA followed by Tukey’s multiple comparisons test. Data presented as mean ± SEM. Statistically significance set at p < 0.05.

## Results

### HIIT training adaptation and exercise performance

We measured numerous parameters including body mass (g), fat percentage (%), lean mass (g), metabolism, and functional tests pre- and post-training for both the SED and HIIT groups. There were no statistical differences in any of the measurements between the groups before training (See **Supplemental Datasets 1-6**).

Initially, the mice ran three training sessions per week for 3 intervals but gradually worked up to 7 intervals--with two-three sessions per week continued as maintenance training during post-training functional testing. The HIIT mice increased in both volume (number of intervals) and intensity (velocity during intervals) as the training study progressed. Maximum running speed (used to determine the interval percentages) significantly increased from the initial test in pre-testing to the mid-trial test, to the post-testing value in HIIT but not in SED (**Figure S1**), which suggests adaptation to aerobic capacity due to training.

### HIIT preserves physical and cognitive function in aged female mice

To determine whether HIIT mitigates age-associated functional decline, the mice tested for physical and cognitive function before and after the training. We measured physical function using rotarod (overall motor function), VWR (volitional exercise rate), grip meter (forelimb strength), inverted cling (four-limb strength/endurance), and treadmill max speed test (aerobic capacity, running speed, endurance), as well as x/y/z plane laser grid monitoring (activity rate). VWR declined significantly following training in both SED (−41%, p=0.004) and HIIT (−42%, p=0.002) mice, with no difference between groups (p=0.991) (**Fig. 2A**). Forelimb grip strength decreased significantly in SED mice (−16%, p=0.002), whereas HIIT mice maintained their strength over the training period (p=0.184), although differences between-groups were not significant (p=0.277) (**Fig. 2B**). Rotarod performance was unchanged in SED mice and exhibited a strong trend toward improvement in HIIT mice (+39%, p=0.078) (**Fig. 2C**). Aerobic capacity significantly improved in HIIT mice (+71%, p=0.002) but did not change in SED, resulting in a significant difference between groups (p=0.002) (**Fig. 2D**). Similarly, inverted cling performance declined substantially in SED mice (−67%, p=0.004) but was preserved in HIIT mice (−29%, p=0.025), also with a significant difference between groups (p=0.018) (**Fig. 2E**). Both groups maintained spontaneous cage activity with no significant differences observed (**Fig. 2F**). A comprehensive breakdown of the statistical analysis is in the **Supplemental Results** section and specifics in **Supplemental Dataset S1. Table 1** lists the more pertinent outcomes. These findings demonstrate that HIIT preserves multiple aspects of physical function during aging, with the strongest effects observed in aerobic capacity and endurance-related measures.

**Figure 2.**
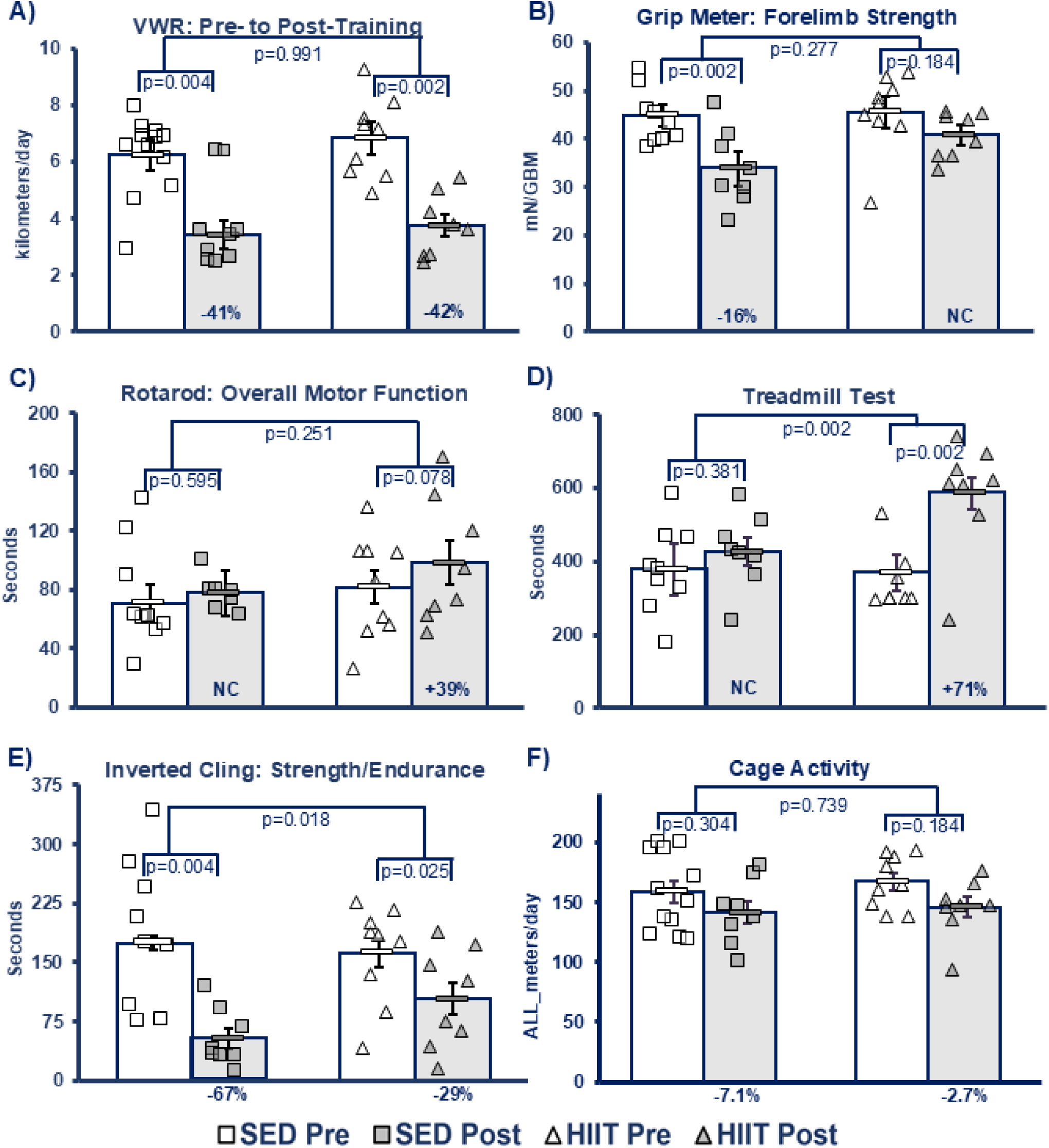
Physical Function A) VWR (voluntary wheel running). Measures volitional exercise rate. **(B) Grip Meter.** Measures strength in the fore limbs. **(C) Rotarod.** Measures overall motor function including elements of balance, coordination, running speed, power production, and endurance. **(D) Treadmill (max speed test).** Measures aerobic capacity and endurance. **(E) Inverted Cling**. Measures whole body strength/endurance. **(F) Cage Activity**. Measures activity in cage via X/Y/Z plane laser motion detectors. **Key:** Open shapes = pre-intervention (Pre), Filled Shapes = post-intervention (post); squares = sedentary group (SED), triangles = HIIT (high intensity interval training) group; % (percent) below graphs = percent change with-in groups from pre- to post-intervention; p-value with-in groups from paired t-test; p-value between groups from independent samples t-test of percent change, red circle = outlier (all statistics in figure run including outlier).

**Table 1.**
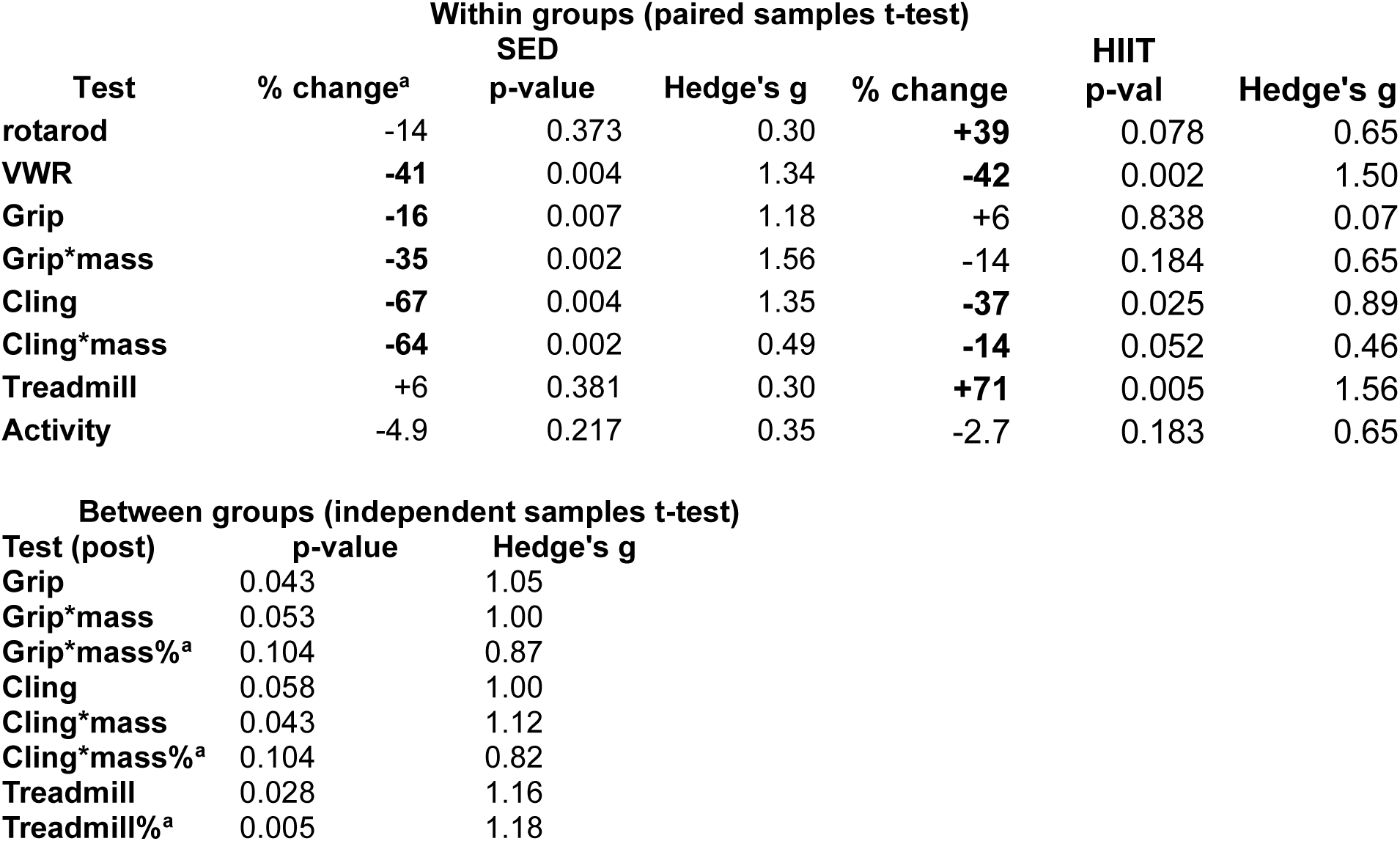
Physical Function Results. For complete results see the online **Supplemental Results** section and **Supplemental Dataset S1**. There were no differences between groups pre-training. **KEY: SED** = sedentary control group, **HIIT =** high intensity interval training group**, bolded** numbers in % (percent) change columns = significant or trend, Pre= before training, Post = after training, NC = no statistical change pre or post, **^a^** = percent change from pre mean to post mean.

HIIT Preserves Function in Older Adult Female Mice
| Within groups (paired samples t-test) |  |  |  |  |  |  |
| --- | --- | --- | --- | --- | --- | --- |
| Test | % change<sup>a</sup> | SED |  | % change | HIIT |  |
|  |  | p-value | Hedge's g |  | p-val | Hedge's g |
| rotarod | -14 | 0.373 | 0.30 | **+39** | 0.078 | 0.65 |
| VWR | **-41** | 0.004 | 1.34 | **-42** | 0.002 | 1.50 |
| Grip | **-16** | 0.007 | 1.18 | +6 | 0.838 | 0.07 |
| Grip*mass | **-35** | 0.002 | 1.56 | -14 | 0.184 | 0.65 |
| Cling | **-67** | 0.004 | 1.35 | **-37** | 0.025 | 0.89 |
| Cling*mass | **-64** | 0.002 | 0.49 | **-14** | 0.052 | 0.46 |
| Treadmill | +6 | 0.381 | 0.30 | **+71** | 0.005 | 1.56 |
| Activity | -4.9 | 0.217 | 0.35 | -2.7 | 0.183 | 0.65 |

**Table 1 Physical Function Results.**
| Between groups (independent samples t-test) |  |  |
| --- | --- | --- |
| Test (post) | p-value | Hedge's g |
| Grip | 0.043 | 1.05 |
| Grip*mass | 0.053 | 1.00 |
| Grip*mass%<sup>a</sup> | 0.104 | 0.87 |
| Cling | 0.058 | 1.00 |
| Cling*mass | 0.043 | 1.12 |
| Cling*mass%<sup>a</sup> | 0.104 | 0.82 |
| Treadmill | 0.028 | 1.16 |
| Treadmill%<sup>a</sup> | 0.005 | 1.18 |

We used the determinants of our composite scoring system, CAB, to measure cognition, which included open field (anxiety and exploratory behavior), NOR (long term memory), y-maze (spatial and working memory), and puzzle-box (memory and executive function). HIIT markedly enhanced executive function in all three measurements (memory, problem solving, and overall), but SED trended to have worse memory, though it did improve in the other two domains to a lesser degree than HIIT. Total puzzle box completion time decreased by 40% in SED mice (p=0.021) and by 73% in HIIT mice (p=0.002), with a significant difference between groups (p=0.046) (**Fig. 3A**). Similarly, performance on the blocked exit component improved significantly only in HIIT mice (−101%, p=0.011) (**Fig. 3B**). NOTE: puzzle box measures show improvements when they decrease. Open field testing revealed increased exploratory behavior following HIIT. Total distance traveled increased significantly in HIIT mice (+89%, p<0.001) compared with SED mice (+75%, p=0.069) (**Fig. 3C**). Time spent in the center of the arena, a measure inversely related to anxiety-like behavior, increased significantly in HIIT mice (+183%, p=0.005) but not SED controls (**Fig. 3D**). Working and spatial memory assessed by spontaneous alternations in the Y-maze improved significantly in HIIT mice (+71%, p=0.002), whereas SED mice showed only a non-significant trend toward improvement (+66%, p=0.077) with an extreme outlier included, but if the outlier is removed there is no change (**Fig. 3E**). In contrast, we observed no differences in novel object recognition discrimination index in either group (**Fig. 3F**), suggesting that *long-term recognition memory* was largely unaffected by exercise, *or* that the test was not sensitive enough to determine differences in this population (see caveats below). See **Table 2** and **Supplemental Dataset S2** for more details.

**Figure 3.**
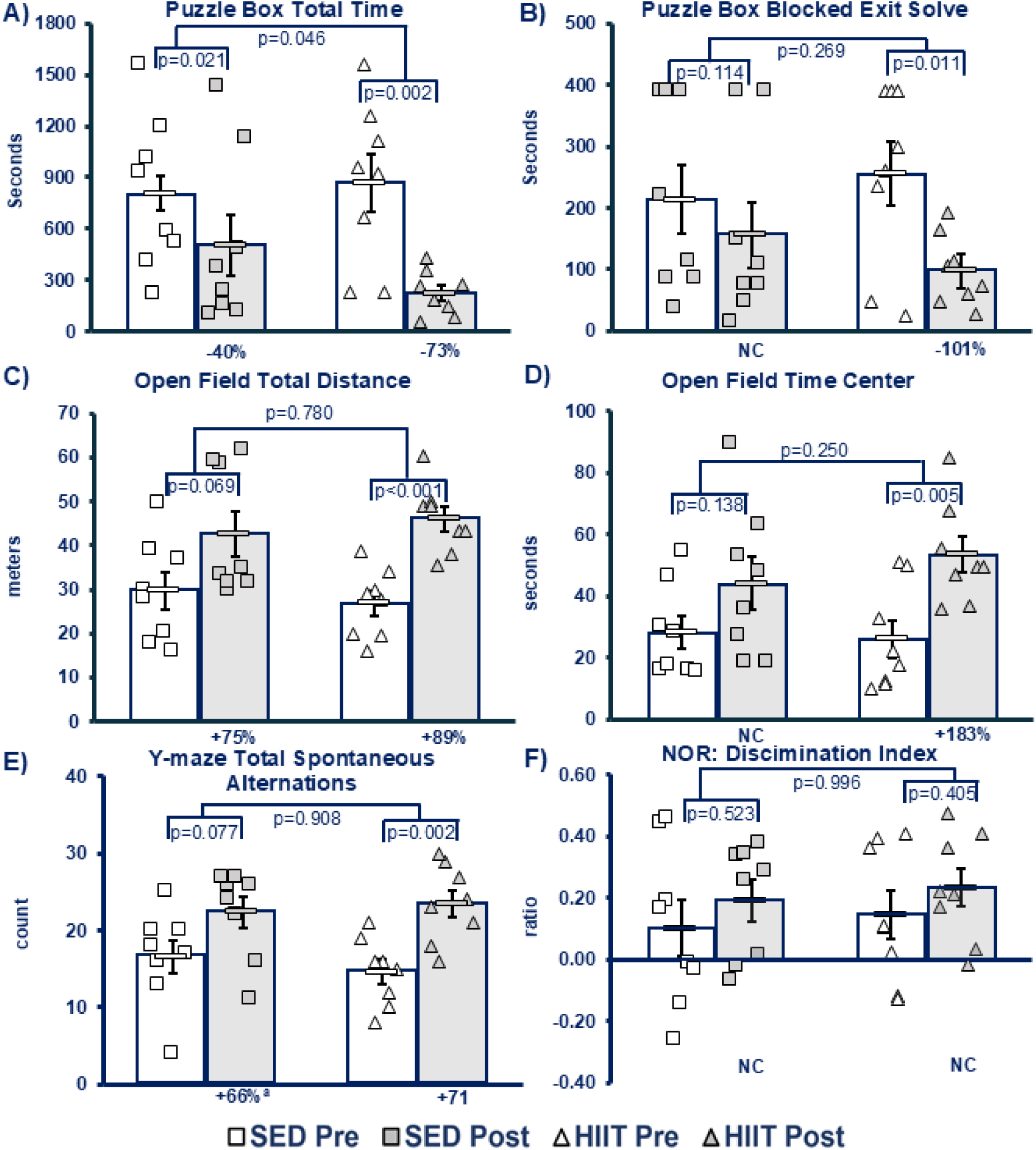
Cognitive Function (A) Puzzle Box Total Time. Measures executive function combined with memory/learning. **(B) Blocked Exit Solve.** Measures executive function. **(C) Open Field Total Distance.** Measures exploratory behavior. **(D) Open Field Time in Center.** Measures anxiety (more time in center of arena = less anxiety). **(E) Y-maze Total Spontaneous Alternations**. Measures exploratory behavior and short-term/spatial memory. **(F) Novel Object Recognition (NOR) Discrimination Index**. Measures long term memory, ratio of (time with novel object – time with familiar object)/(sum of both times). **Key:** Open shapes = pre-intervention (Pre), Filled Shapes = post-intervention (post); squares = sedentary group (SED), triangles = HIIT (high intensity interval training) group; % (percent) below graphs = percent change with-in groups from pre- to post-intervention; NC = no change; p-value with-in groups from paired t-test; p-value between groups from independent samples t-test of percent change, however, in panel **(F)** the p-value between groups independent samples t-test of difference from pre to post. **^a^** There was an extreme outlier in the SED group (+300% SA pre to post, removing this outlier within-group resulted in *no change*, p=0.293; between groups p=0.142). NOTE: Puzzle box scores improved if they decreased.

**Table 2.** Cognitive Function Results. For complete results see the online **Supplemental Results** section and **Supplemental Dataset S2**. **KEY:** NOR = novel object recognition, NO = novel object, discrimination = ratio of time spent on novel object compared to total explorations, SA = total number of spontaneous alternations, SA% = percentage of SA of total arm entries, PB1_2 = first test of puzzle box, PB3tot= second test of puzzle box, Pbtot = puzzle box total score (PB1_2 + PB3tot), CAB = cognitive assessment battery (compound scoring system), SED = sedentary control group, HIIT = high intensity interval training group**, bolded** numbers in % (percent) change columns = significant or trend, Pre= before training, Post = after training, NC = no statistical change pre or post between groups, **^a^** = percent change from pre mean to post mean (larger number = better in open field, NOR, and y-maze, but lower number = better in puzzle box).

| Test | Parameter | % change | SED |  | % change | HIIT |  |
| --- | --- | --- | --- | --- | --- | --- | --- |
|  |  |  | p-val | Hedge's g |  | p-val | Hedge's g |
| Open Field | total distance | -14 | 0.373 | 0.30 | **+39** | **0.078** | 0.65 |
|  | center time | +57 | 0.163 | 0.49 | **+105** | **0.013** | 1.03 |
| NOR | discrimination | +90 | 0.523 | 0.42 | +36 | 0.632 | 0.16 |
|  | NO time | **+51** | **0.053** | 0.73 | **+300** | **0.096** | 0.53 |
|  | total exploration | +31 | 0.239 | 0.26 | +24 | 0.154 | 0.20 |
| Y-maze | SA | **+35** | **0.060** | 1.40 | **+61** | **0.001** | 1.63 |
|  | SA% | +11 | 0.329 | 0.33 | +9 | 0.163 | 0.49 |
| Puzzle Box | PB1_2 | **-42** | **0.015** | 1.01 | **-77** | **0.004** | 1.33 |
|  | PB3tot | **+32** | **0.065** | 0.69 | **-72** | **0.005** | 1.25 |
|  | Pbtot | **-37** | **0.021** | 0.93 | **-74** | **0.002** | 1.50 |

**Table 2 Cognitive Function Results.**
| Test | Parameter | p-val | Hedge's g |
| --- | --- | --- | --- |
| Puzzle Box | PB_%change | **0.046** | 1.22 |

To determine whether these HIIT-induced adaptations translated into preservation of overall function, we calculated the composite physical (CFAB) and cognitive (CAB) scores. SED mice exhibited a significant decline in overall physical function over the training period (p=0.001), whereas HIIT mice maintained physical performance (p=0.881) (**Fig. 4A**). Consistent with this, HIIT preserved the CFAB score over the time period while SED declined (ΔCFAB: HIIT -0.772, no change, vs. SED -4.25, p=0.021) (**Fig. 4B**). Cognitive composite scoring revealed that SED mice exhibited no overall improvement, but HIIT increased CAB scores by 34% (p=0.022) (**Fig. 4C**), with significantly greater changes observed in HIIT mice compared with SED controls (p=0.039) (**Fig. 4D**). Collectively, these findings demonstrate that HIIT preserves overall physical function and improves cognitive performance in aged female mice. See **Table 3** for details on pertinent outcomes.

**Figure 4.**
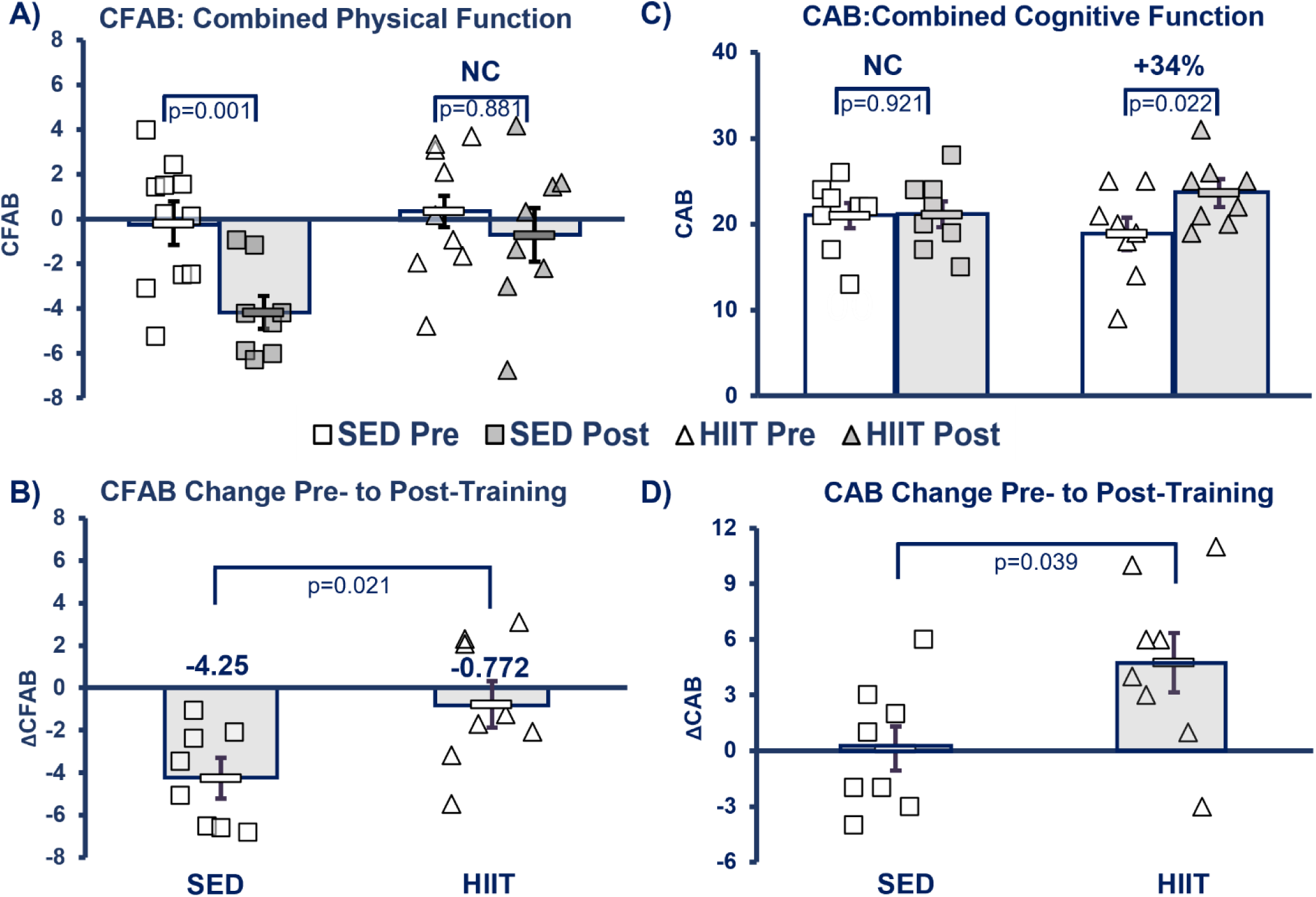
CFAB and CAB Composite Scores (A) CFAB (Comprehensive Functional Assessment Battery); (B) Change in CFAB (ΔCFAB); (C) CAB (Cognitive Assessment Battery); (D) Change in CAB (ΔCAB). Key: Open shapes = pre-intervention (Pre), Filled Shapes = post-intervention (post); squares = sedentary group (SED), triangles = HIIT (high intensity interval training) group; % (percent) below graphs = percent change with-in groups from pre- to post-intervention; p-value with-in groups from paired t-test; p-value between groups from independent samples t-test of percent change.

**Table 3.**
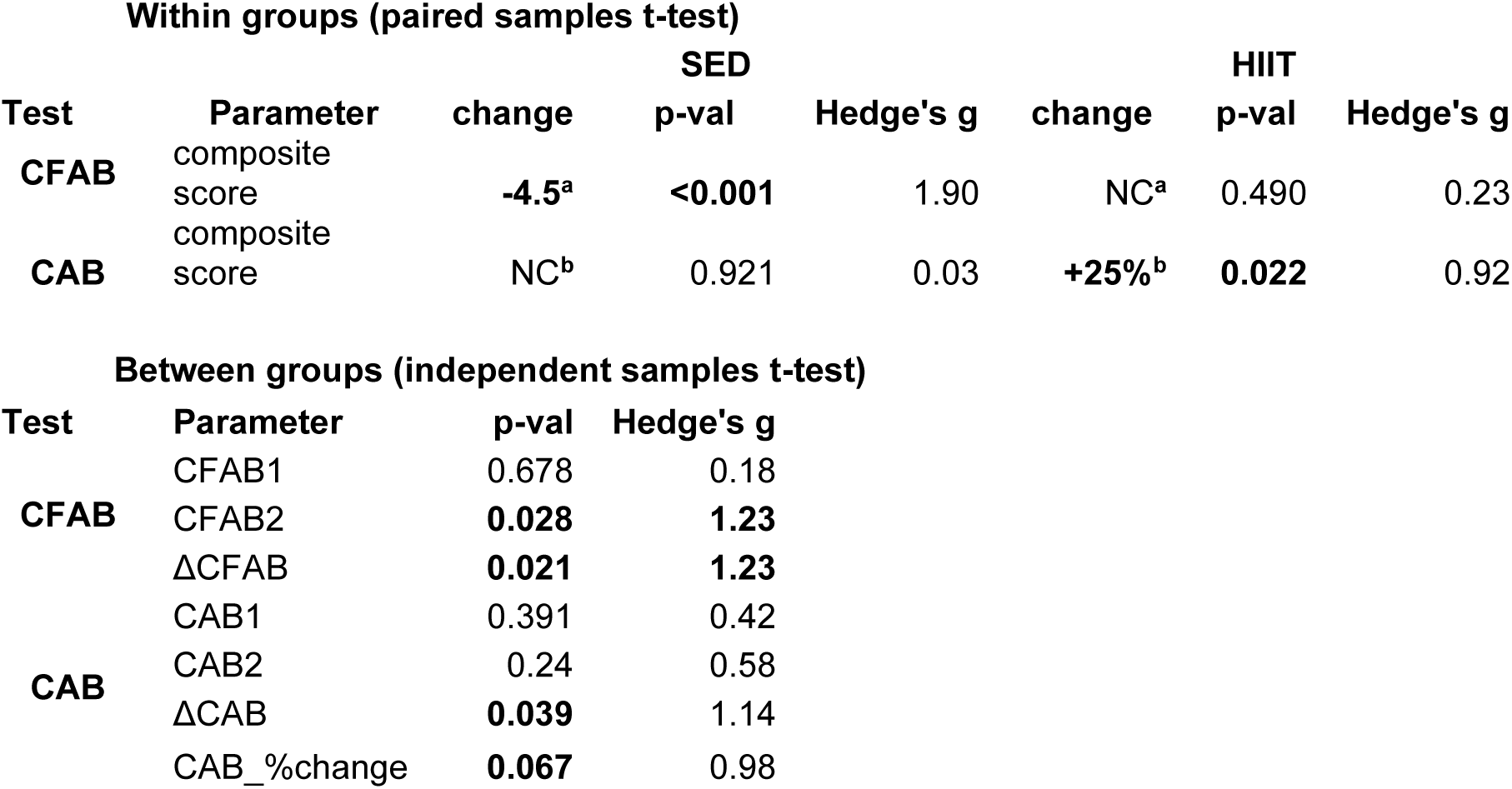
Composite Scoring System Results: CFAB and CAB. For complete results see the online **Supplemental Results** section and **Supplemental Dataset S2 and S3**. **KEY:** CFAB = comprehensive functional assessment battery for physical function, CAB = cognitive assessment battery (compound scoring system), SED = sedentary control group, HIIT = high intensity interval training group**, bolded** numbers = significant or trend, Pre= before training, Post = after training, NC = no statistical change pre or post between groups, **^a^**= difference of post-CFAB – pre-CFAB (more negative = worse outcome), **^b^**= percent change from pre mean to post mean (larger number = better).

### HIIT delays frailty progression in aged female mice

Because preservation of functional capacity is closely linked to frailty status, we assessed frailty scores in the post-training SED and HIIT groups using a modified version of our previously published reverse translation of the Fried Frailty Phenotype.(20,21,23,29) Prior to training, none of the mice in either group were frail, but one mouse in the HIIT group was pre-frail. In the SED group, frailty status worsened over the training period, with two mice becoming pre-frail and four mice progressing to frailty, leaving only two animals classified as non-frail at study completion (**Fig. 5A**). In contrast, HIIT prevented frailty progression. Four HIIT mice remained non-frail and four were pre-frail following the training, with *no animals* progressing to the frail category. To determine whether overall physical function reflected frailty status, we compared post-training CFAB scores with frailty scores (**Fig. 5B**). Lower CFAB scores were strongly associated with greater frailty burden, with all pre-frail mice exhibiting CFAB scores below -2 and all frail mice exhibiting CFAB scores below -6. These findings indicate that reductions in overall physical function closely track frailty progression. These results suggest that HIIT initiated late in life can mitigate frailty development and onset.

**Figure 5.**
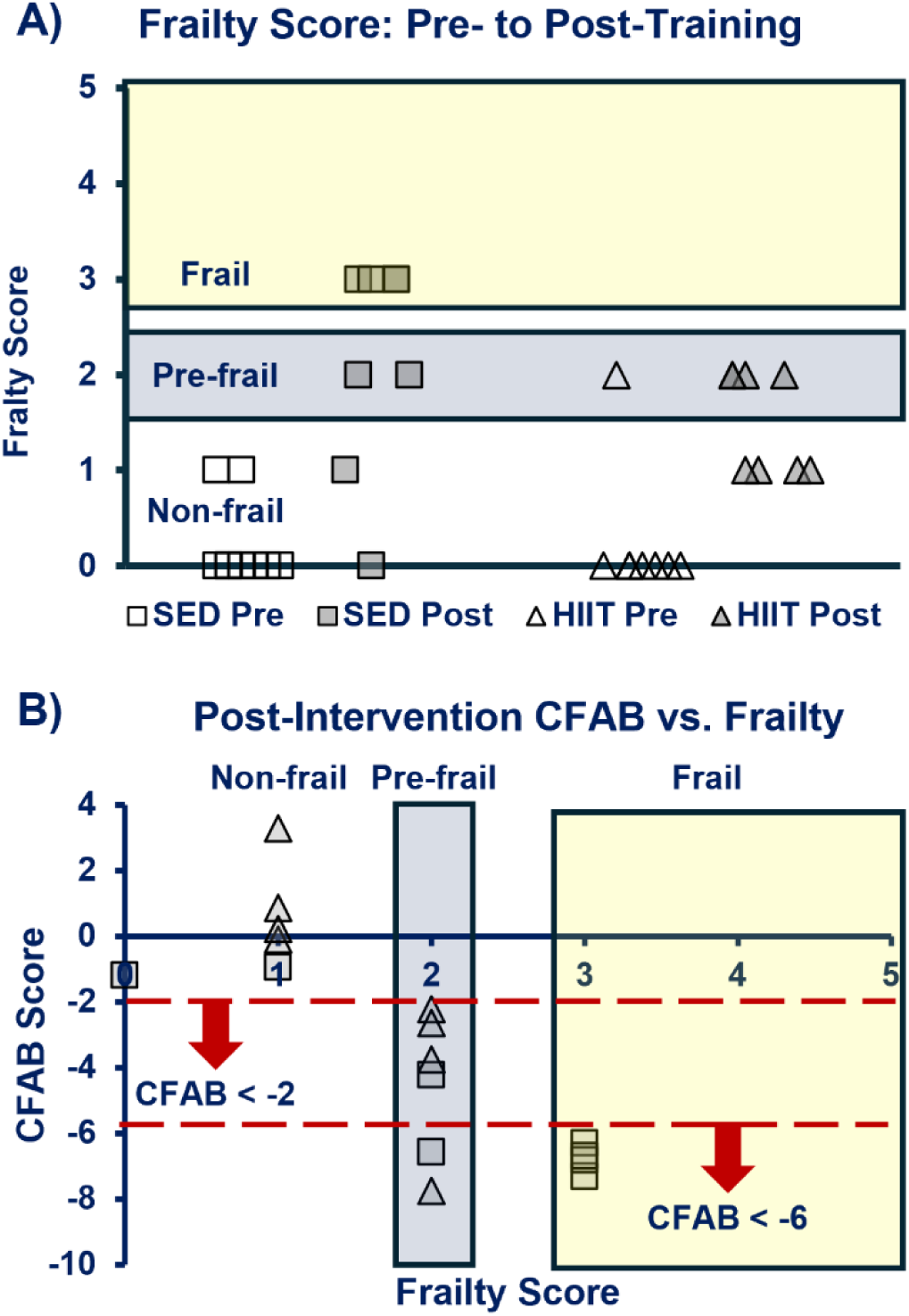
Frailty. A) Frailty Score Pre- to Post-Intervention. Frailty score ≥ 3 = frail, 2 = pre-frail, 1 or 0 = non-frail. B) **CFAB (Comprehensive Functional Assessment Battery) versus frailty score.** CFAB measures overall physical function with a lower score equaling worse functional ability. NOTE: All pre-frail mice scored less than -2 in CFAB, and all frail mice scored less than -6 in CFAB indicating CFAB serves as a proxy for frailty measurements**. Key:** Open shapes = pre-intervention (Pre), Filled Shapes = post-intervention (post); squares = sedentary group (SED), triangles = HIIT (high intensity interval training) group. Dotted lines indicate CFAB cutoffs for prefrail (−2) and frail (−6).

### HIIT improves cardiovascular health and alters metabolic function during aging

To determine whether the functional benefits of HIIT extended to cardiovascular and metabolic health, we evaluated blood pressure, cardiac structure, body composition, and whole-body metabolism. HIIT significantly reduced systolic, diastolic, and mean arterial blood pressure following training, whereas SED mice exhibited no significant changes (**Fig. 6**). Systolic blood pressure decreased by 12.5% (p=0.003), diastolic blood pressure decreased by 15% (p=0.004), and mean arterial pressure decreased by 14% (p=0.004) in HIIT mice. Although between-group comparisons did not reach statistical significance, we observed large effect sizes for changes in systolic (g=0.85), diastolic (g=0.75), and mean arterial pressure (g=0.73).

**Figure 6.**
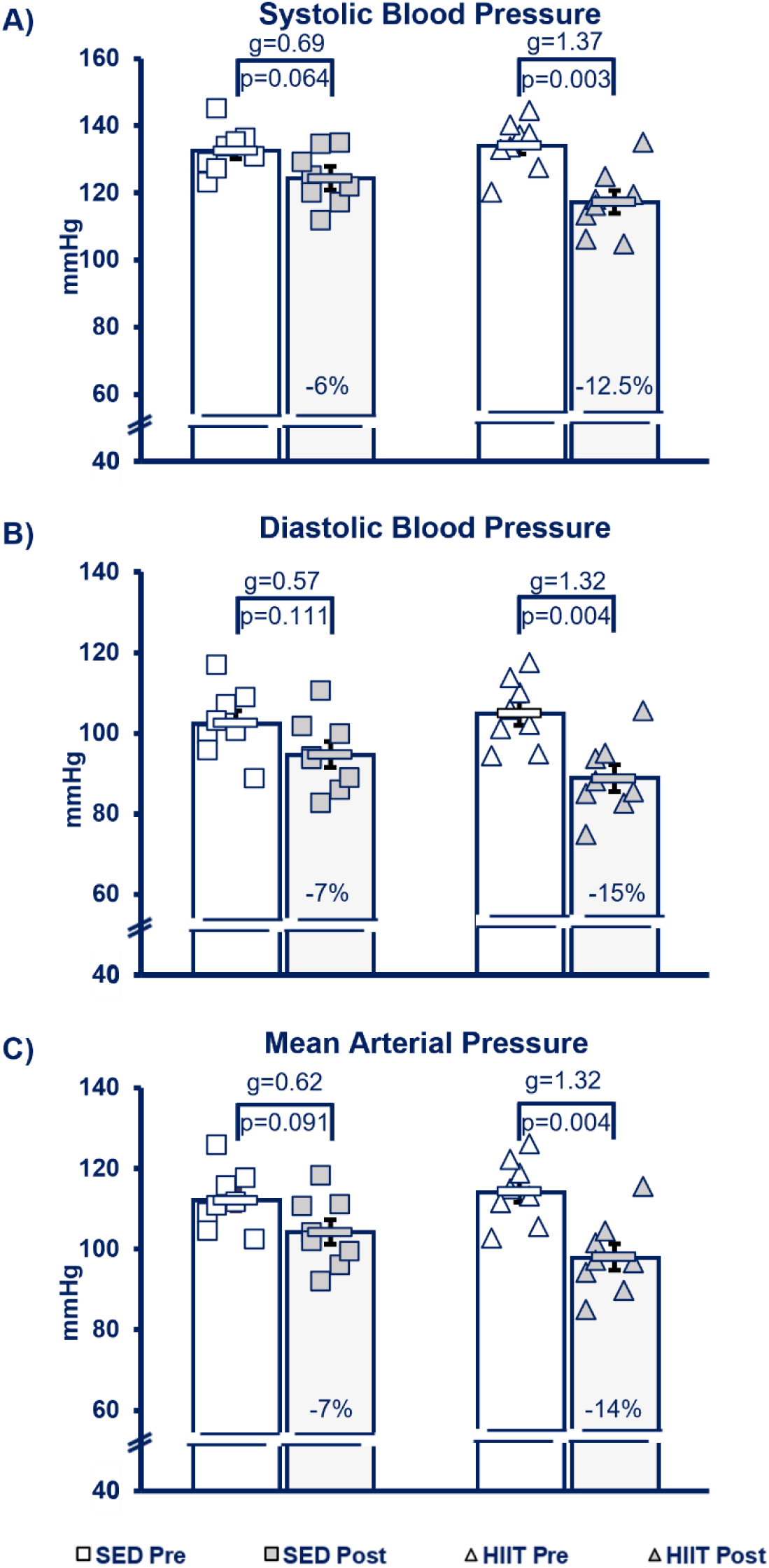
Blood Pressure. A) Systolic Blood Pressure, B) Diastolic Blood Pressure, C) Mean Arterial Pressure. Key: Open shapes = pre-intervention (Pre), Filled Shapes = post-intervention (post); squares = sedentary group (SED), triangles = HIIT (high intensity interval training) group; % (percent) below graphs = percent change with-in groups from pre- to post-intervention; p-value with-in groups from paired t-test; p-value between groups from independent samples t-test of percent change.

Echocardiography revealed no statistically significant differences in cardiac structural parameters between groups (**Supplemental Dataset 3**). However, HIIT mice exhibited trends toward reduced ventricular wall thickness and attenuation of age-associated cardiac remodeling. Left ventricular anterior wall thickness in systole (LVAWd) reduced by approximately 9% in HIIT mice (g=0.72), whereas SED mice exhibited a trend toward increased wall thickness (+9%; g=0.60), suggesting that exercise may limit the progression of adverse cardiac structural remodeling.

We weighed the mice monthly during aging up to training and then weekly once training commenced to track body mass. Body composition analysis demonstrated that both groups gained body weight, lean mass, and fat mass over the training period. We determined no significant differences between SED and HIIT mice for body mass, lean mass, fat mass, or body fat percentage before or after training (**Fig. S2** and **Supplemental Dataset S4**). Although HIIT mice exhibited a significant increase in body fat percentage (+6.6%, p=0.006), overall body composition remained comparable between groups, indicating that HIIT-induced functional improvements occurred despite limited effects on overall body composition.

Despite minimal changes in body composition, HIIT training resulted in several metabolic adaptations. Pre- and post-training the mice spent 7 days in Promethion Metabolic Cages. Both groups exhibited similar reductions in carbon dioxide production and respiratory exchange ratio following training. However, HIIT mice demonstrated significant decreases in oxygen consumption, maximal carbon dioxide production, caloric expenditure, and food intake, while SED did not (**Fig. S3** and **Supplemental Dataset S5**). Collectively, these findings suggest that HIIT alters whole-body metabolic responses during aging, consistent with improved metabolic efficiency and adaptations in substrate utilization.

### HIIT modulates neuroinflammatory markers in the aged brain

To determine whether HIIT modifies neuroinflammatory phenotypes in aged female mice, we used immunofluorescence to assess glial activation and inflammatory cytokine expression in the hippocampus. GFAP expression and morphological complexity assessed astrocytic activation. Although aged mice had elevated GFAP expression relative to young controls (p<0.05), HIIT did not significantly alter GFAP expression compared with SED aged mice (p=0.165) (**Fig. 7A,B**). However, Sholl analysis revealed a trend toward reduced astrocytic ramification in HIIT mice compared with SED mice (p=0.076), suggesting that exercise may partially attenuate age-associated astrocyte hypertrophy (**Fig. 7C**). We next examined microglial activation to determine whether HIIT influences innate immune responses in the aged brain. IBA1 expression increased in SED aged mice compared with young controls (p<0.05). HIIT reduced IBA1 expression relative to SED mice, although this reduction did not reach statistical significance (p=0.081), and expression remained elevated compared with young mice (p=0.061) (**Fig. 7D,E**). Morphological analysis using skeletonization revealed significantly greater branching in young mice compared with both aged groups (p<0.05), consistent with age-related microglial activation. Importantly, HIIT mice exhibited a trend toward increased microglial branching compared with SED mice (p=0.054), indicating a partial normalization of microglial morphology with exercise (**Fig. 7F**). IL-1β expression quantification determined whether these glial phenotypic changes were associated with alterations in inflammatory signaling. Both HIIT and SED aged mice exhibited increased IL-1β compared with young controls (p<0.05). Notably, HIIT significantly reduced IL-1β expression compared with SED aged mice (p<0.05), indicating that exercise attenuates neuroinflammatory cytokine expression in aging (**Fig. 7G**). Collectively, these findings indicate that HIIT partially reveres age-associated neuroinflammatory phenotypes.

**Figure 7.**
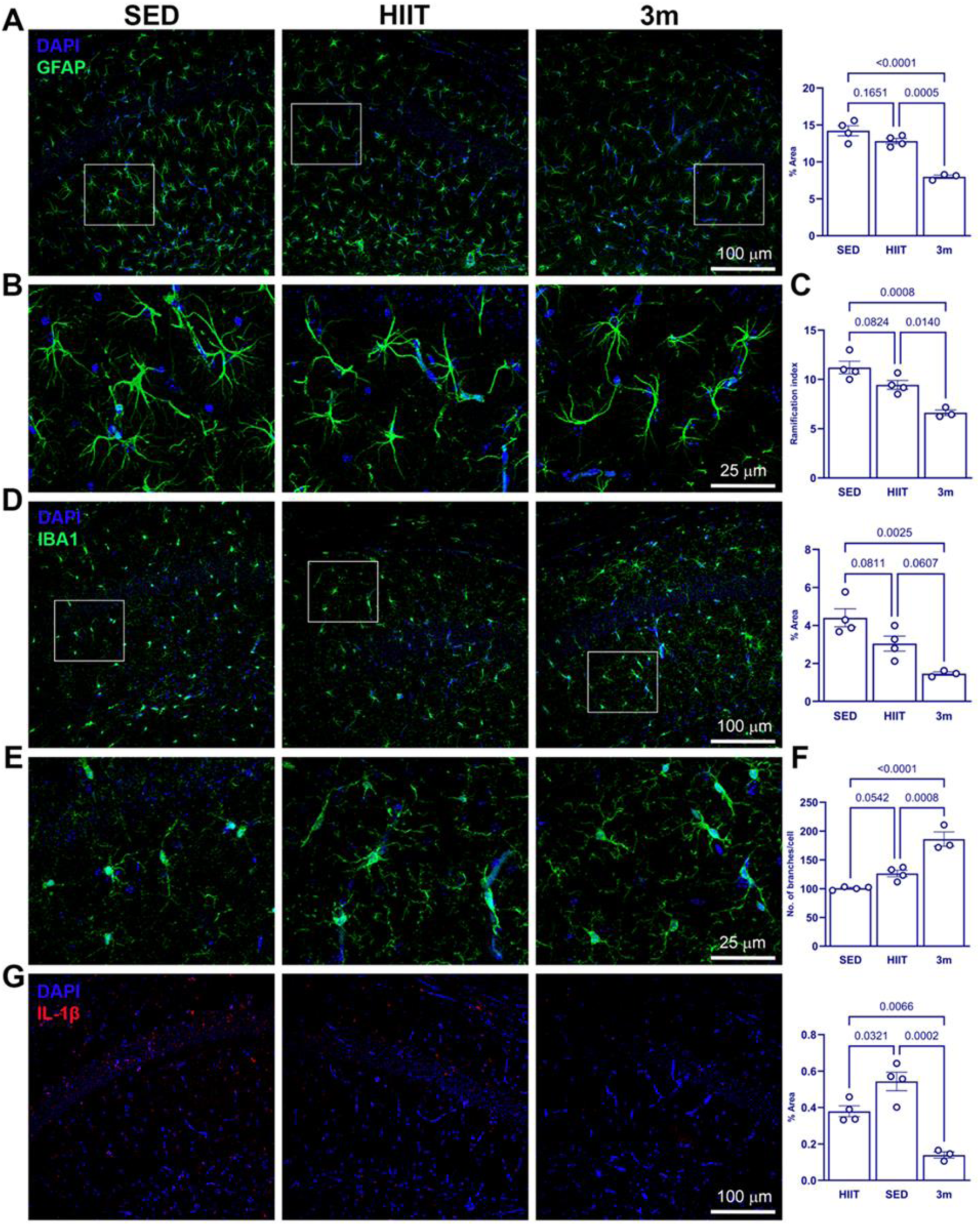
HIIT partially attenuates neuroinflammatory signaling and glial activation in aged mice. **(A)** Representative images and quantification of GFAP expression demonstrate increased astrocytic activation in aged mice compared with young controls, with no significant difference between HIIT and SED aged mice. **(B)** Higher-magnification images illustrate astrocyte morphology across groups. **(C)** Sholl-based ramification index reveals increased astrocytic complexity in aged mice, with a trend toward reduced ramification in HIIT compared with SED mice. **(D)** Representative images and quantification of IBA1 expression show increased microglial activation in SED aged mice, with partial reduction following HIIT. **(E)** Higher-magnification images highlight microglial morphological differences between groups. **(F)** Skeletonization analysis demonstrates reduced microglial branching in aged mice, with a trend toward increased branching in HIIT mice relative to SED mice. **(G)** IL-1β expression increased in aged mice and significantly reduced in HIIT compared with SED controls. Data presented as mean±SEM. Data analyzed using one-way ANOVA followed by Tukey’s multiple comparisons test. n=10 cells/image, 5 images/mouse, 4 mice/group. Statistical significance was set at p<0.05.

### HIIT influences cardiac remodeling markers in aged mice

To determine whether HIIT influences structural remodeling of the aging heart, we assessed markers of cardiomyocyte hypertrophy, fibroblast activation, and extracellular matrix remodeling in the left ventricle. Cardiomyocyte cross-sectional area did not differ significantly among groups; however, SED aged mice demonstrated a trend toward cardiomyocyte hypertrophy compared with young mice (p=0.077). We did not observe this trend in HIIT mice, suggesting that exercise may limit age-associated cardiomyocyte hypertrophy (**Fig. 8A**). To determine fibroblast activation, we measured αSMA expression. SED aged mice exhibited significantly increased αSMA expression compared with young controls (p<0.05), whereas HIIT mice showed only a modest, non-significant increase relative to young mice (p=0.088). However, HIIT and SED had no significant difference (p=0.386), indicating that HIIT did not significantly alter myofibroblast differentiation at the time point assessed (**Fig. 8B**). Expression of the gap junction protein Cx43 was not significantly different between groups, suggesting that HIIT did not substantially alter myocardial gap junctions (**Fig. 8C**). Picrosirius red staining quantified collagen deposition, showing aged SED mice demonstrated significantly greater collagen accumulation compared to young controls (p<0.05), consistent with increased fibrotic remodeling. In contrast, collagen deposition in HIIT mice reduced relative to SED aged mice and did not significantly differ from young controls, suggesting HIIT attenuated age-associated extracellular matrix remodeling and fibrosis (**Fig. 8D**). Taken together, these results suggest that HIIT attenuates key features of age-associated cardiac remodeling.

**Figure 8.**
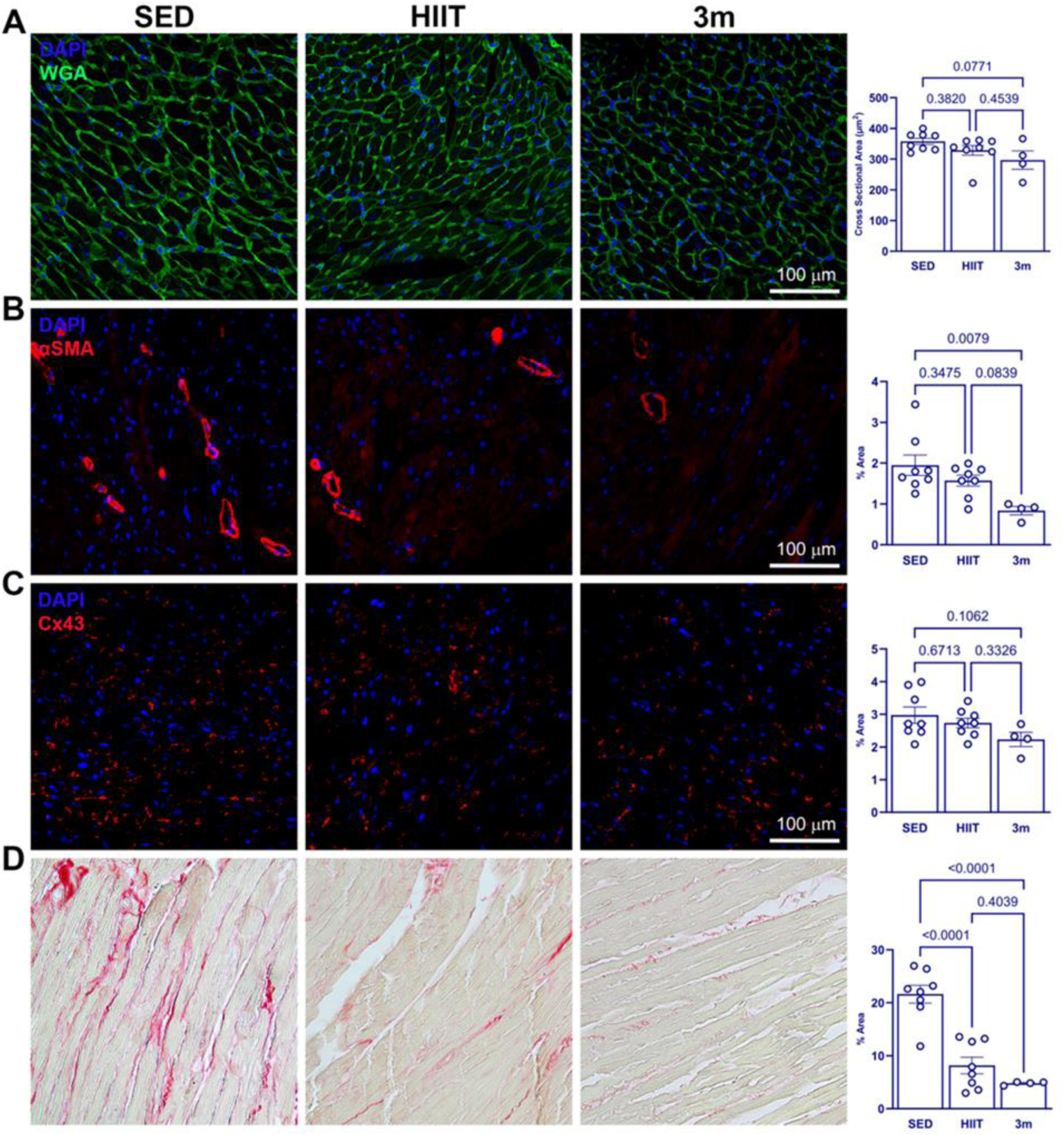
HIIT modulates markers of cardiac remodeling in aged mice. **(A)** Quantification of cardiomyocyte cross sectional area reveals a trend toward hypertrophy in SED aged mice not observed in HIIT mice. **(B)** αSMA expression significantly increased in SED aged mice compared to young controls, with a smaller, non-significant increase in HIIT mice. **(C)** Cx43 expression does not differ among groups. **(D)** Picrosirius red staining and quantification of collagen content. Data presented as mean±SEM. Data analyzed using one-way ANOVA followed by Tukey’s multiple comparisons test. For panel A: n=5 cardiomyocytes/image, 4-5 images/group; for panels B-D: n=4-5 images/group; 8 mice/group (aged), 4 mice/group (young). Statistical significance was set at p<0.05.

### HIIT alters skeletal muscle fiber composition in aged female mice

To determine whether HIIT alters skeletal muscle phenotype during aging, we measured soleus muscle fiber composition and morphology using myosin heavy chain (MHC) isoform immunofluorescent staining. There were alterations in type I (MHC1) and type IIa (MHC2a) fibers distribution between SED and HIIT mice (**Fig. 9A,B**). HIIT mice exhibited more type I fibers and a corresponding reduction in type IIa fibers compared to SED. Type I fibers increased from 57% in SED mice to 64% in HIIT mice, whereas type IIa fibers decreased from 43% to 36%. Quantification of myofiber CSA revealed no statistically significant differences between groups for either fiber type (**Fig. 9C**). However, type IIa fibers exhibited a large effect size, with HIIT demonstrating a 47% greater mean CSA than SED controls (g=0.867), suggesting a potential effect that may not have reached significance due to large individual variation. Analysis of fiber type distribution revealed significant alterations in soleus muscle composition following HIIT (**Fig. 9D**). See **Supplemental Dataset S6** for further details. Overall, these findings suggest that HIIT promotes remodeling of soleus muscle fiber composition during aging in female mice.

**Figure 9.**
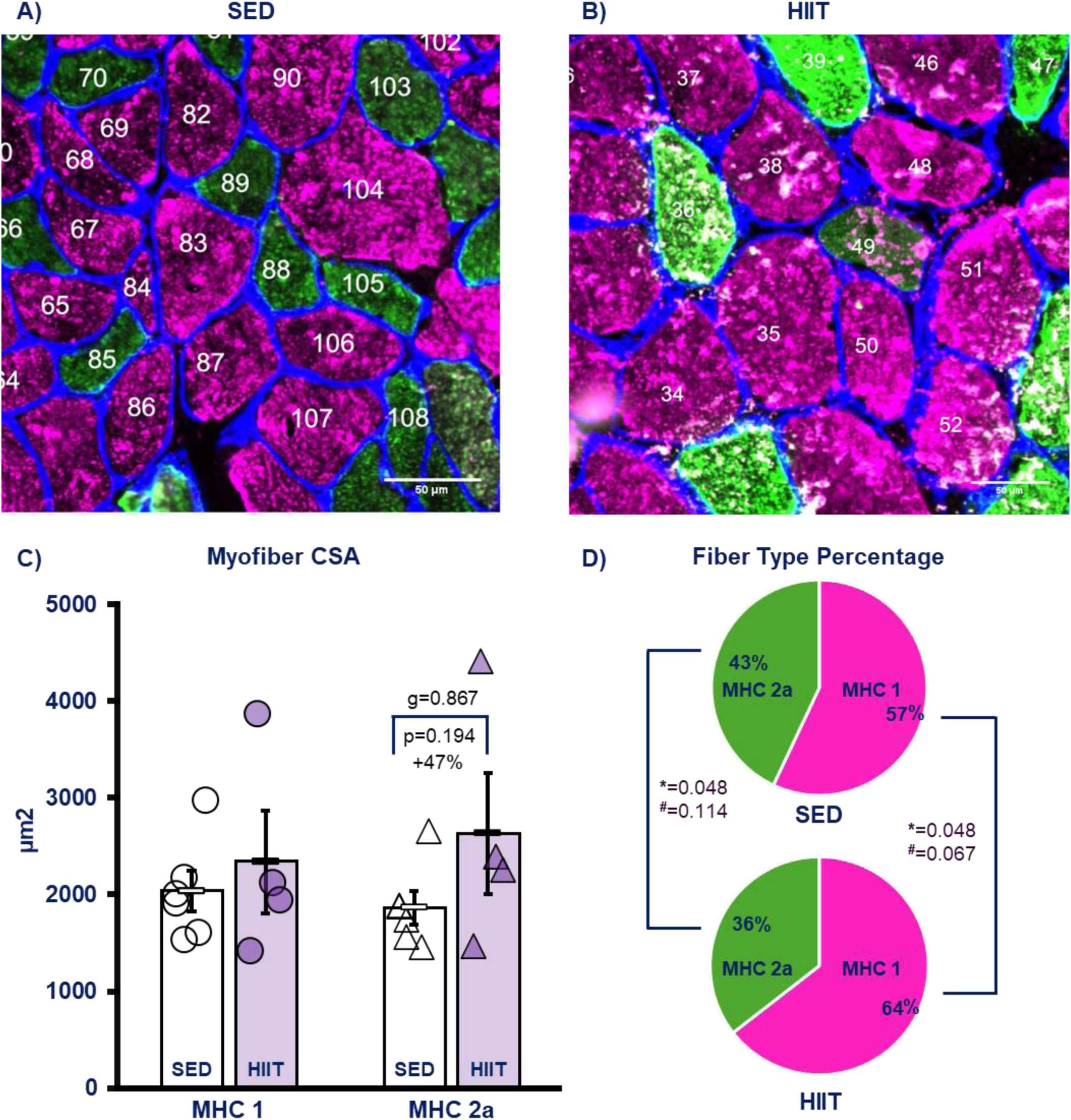
Soleus Muscle Morphology A) SED Representative Image, B) HIIT Representative Image. KEY: Pink = myosin heavy chain type 1 (MHC 1) and green = myosin heavy chain type 2a (MHC 2a), Green + Pink = hybrid 2a/1 (note cell # 49 in Panel B), blue = laminin**. C) Cross-sectional Area (CSA).** No significant differences but strong Hedge’s g effect size in MHC 2a. **D) Fiber Type Percentage.** There were significant differences between groups (non-parametric via **^#^**Mann-Whitney U-test and *Median Test) with HIIT having fewer 2a fibers and more type 1. **KEY:** SED = sedentary (open symbols), HIIT = high intensity interval training (filed symbols), MHC 1 = circles, MHC 2a = triangles.

## Discussion

The primary purposes of this study were to extend our prior work done in male mice to include aged females, to determine the effect of HIIT to modulate frailty, age-associated dysfunction in physical, cognitive, and cardiovascular domains, uncover how HIIT impacts brain, heart, and muscle tissues, and reveal potential mechanistic clues. To our knowledge this is the first such comprehensive evaluation of HIIT on older adult female mice. We hypothesized that HIIT would modulate age-related decline in all the systems we investigated and would either improve or preserve functional capacity. Our prior work in male mice demonstrated HIIT could preserve physical function during aging, and now we have evidence in females of not only physical function, as we expected, but also other functional domains. We uncovered evidence leading us to conclude HIIT in older female promotes improved cardiovascular and cognitive health. Much of the previous literature has previously focused on males, so the addition of female investigation is critical to the field.(16)

### Body Composition

While cardiovascular outcomes from endurance training have been widely observed in different populations, other adaptations such as body composition, physical function, cognitive function, and various molecular and cellular changes are often more nuanced.(30) For example, our lab has observed discrepancies between the sexes and ages in body composition changes due to endurance training, nuanced further by exercise type. With lower volume HIIT or voluntary wheel running (VWR), adult exercised males (10m at study end) maintained both body mass and fat%, whereas their sedentary counterparts gain a large amount in both.(5) In contrast, the same study showed the older adult males significantly losing both fat% and body mass, though interestingly the VWR group lost more fat. The sedentary old males maintained both mass and fat%. Recently, we studied lower volume HIIT on middle-aged male mice (17m at study end) and found that both sedentary and HIIT groups increased in body mass and fat%, though lean mass tended to be elevated in the HIIT mice with a strong effect size (0.746).(14) However, using a different exercise paradigm (power training—essentially HIIT while wearing a weighted vest), older male mice did lower fat% post-training.(31)

In the current study, the older female mice gained weight in both groups (SED and HIIT). Body fat also increased in both groups but reached statistical significance only in HIIT group (large effect size in both groups). These findings were recently echoed by the MoTrPAC research group using a moderate intensity continuous training (MICT) rat model, in which after 8 weeks, adult male rats lost 5% body fat and body weight, whereas female rats did not lose body fat and instead gained weight.(32) During aging we have shown longitudinally that body mass changes in sedentary males and sedentary females do not follow the same patterns, with males gaining weight from 6-18 months and then stabilizing body mass from 18-29m, while females gain large amounts of body mass from 6-24m, and then lose mass from 24-28m.(17) The same study demonstrated female mice gaining large amount of fat between adulthood and middle-age, but then losing fat as they progressed from older adult status to very advanced age. Seldeen and colleagues have also extensively studied HIIT training in mice, noting that in both older males and females there was a significant, though modest, decrease in body weight for both HIIT and SED.(6,7,33) Aerobic exercise in people shows some limited benefit for weight loss and body composition change at low volume/intensity, but can become clinically significant with greater volume and intensity (>150 minute/week at moderate or greater intensity).(34) In humans, there is evidence that HIIT improves body composition in adults, but as in our preclinical studies, there are nuances based on population studied, HIIT exercise type (e.g., cycling versus running), and protocols used.(35) Together, these findings suggest that the effects of exercise training on body mass and adiposity are sex-, age-, and protocol-dependent, and that HIIT in older female mice may improve functional outcomes without necessarily preventing age-associated increases in body weight or fat mass. Is this correct?

### Physical Function, Metabolism, and Frailty

We inevitably lose some percentage of physical function and exercise capacity with advancing age. Preserving, maintaining, or improving physical function during aging is critical for older adults to be able to perform activities of daily living, maintain independence, and prevent the onset of disability. One promising intervention to preserve this function is exercise.

We have previously well-documented the age-related decline of physical and contractile function in mice, including sexual dimorphisms in the patterns of functional loss.(17,22,36,37) Other labs have observed similar findings in humans.(38) Using exercise as a countermeasure to slow the trajectory of this functional decline is a viable intervention strategy.(5,31,39) We previously demonstrated the efficacy of HIIT to preserve physical function in aging male mice.(5) In our current study, we determined that HIIT reduced age-associated loss of physical function in older female mice.

Resistance training is often touted as a defense against the onset of sarcopenia (age-related loss of muscle mass and strength that contributes to a decline in physical function). Resistance training can increase or preserve muscle mass and strength even in older and frail individuals.(40) In general, muscle force is dictated by the number of concurrent myofilament contracting, hence the maxim that more and larger myofibers = more muscle force (the bigger the muscle the greater the potential for force development). However, loss of *muscle power* predates and exceeds the loss of muscle force during aging, particularly at tasks closer to maximal force output.(37,41) Previously we determined that power training (essentially a HIIT protocol while wearing a weight pack) improved power output in older and adult mice at high percentages of maximum force (60%, 80% and 90%).(31)

While our current HIIT study did not directly measure power, we do note that the fiber type percentages altered significantly with HIIT training favoring a myosin heavy chain (MHC) type 1 distribution in the soleus. Type 2a muscle fibers (fast twitch, aerobic/glycolytic) produce, in general, *more force at a greater contractile velocity* than type 1 fibers (more predominant in our SED mice). While the fiber type distribution changed with HIIT favoring MHC1, there was some evidence (strong effect size, Hedge’s g=0.867) that the remaining type 2a muscles grew substantially (+47%) in size which may compensate by producing greater force (larger cross-sectional area) from fewer myofibers, resulting in no loss of, or even a gain of power. Type 2a and type 1 muscle fibers are both fatigue resistant, though type 1 may be better at aerobic activity, so we hypothesize that this SOL adaptation to HIIT overall may be expected to change muscle endurance without effecting power negatively. An additional parameter of muscle regulation of power production involves the phosphorylation of the regulatory myosin light chain (MLC), which exists as multiple isoforms that can mix within myofibers and cause a difference in power production among fibers with the same overall MHC isoform. In prior work we found that increasing MLC 3f in older adult rats prevented age-related loss of contractile velocity, but to our knowledge the effect of HIIT on MLC isoforms is unknown.(42) Thus, while outside the scope of the current study, more future research is needed to confirm this new hypothesis, including direct measurement of contractile velocity and force output *ex vivo* to compare SED versus HIIT power production, determining regulatory MLC composition, and assessment of fiber type composition and myofiber CSA in other locomotor muscles, such as the gastrocnemius and quadriceps.

Muscle metabolism is also influenced by exercise. At rest, individuals with greater muscle mass use more calories (higher basal metabolic rate) than those who are more obese since adipocytes are relatively metabolically inactive compared to myofibers. Muscle, being a highly plastic tissue, adapts to demands, for example by hypertrophy after resistance training or increasing the number of mitochondria after endurance training. While we did not measure metabolic processes *in vitro* in the current study, it is interesting to note that both SED and HIIT mice saw a similar reduction in RER while having similar activity levels and weight/fat gains, though energy expenditure (Kcals/day) and food intake by HIIT groups was markedly reduced, suggesting improved metabolic efficiency by the exercise group. The drop in food intake by HIIT during their week in the metabolic chambers, where they did not have training sessions and maintained similar activity and volitional exercise rates as the SED, also suggests that they may have been consuming more food during the time spent training to offset the additional calories burned, resulting in the body composition results of our group of older females. There is some evidence that exercise alone has been shown to be relatively modestly effective at altering body composition or body weight in healthy adults, but does have a synergistic effect when combined with a dietary intervention.(30,43)

In prior work, we and others have found that exercise is sufficient to prevent or ameliorate the onset of frailty in mouse models.(6,7,21) In the current work, sedentary older adult female mice progressed towards frailty while that progression was retarded with HIIT. Importantly, we confirmed that our CFAB score can serve as a proxy for the frailty phenotype system we originally reverse translated from the human version published by Fried and colleagues.(29) In humans similar results can be expected.(44)

### Cognitive Function

Physical activity (PA) and exercise may play a key role in risk reduction for the onset of Alzheimer’s Disease and related dementias (ADRD) onset and/or slowing disease progression while also mitigating age-related cognitive decline in general.(45) Increasing aerobic fitness is an intervention to reduce age-related cognitive decline and potentially the onset of mild cognitive impairment (MCI) or ADRD via numerous mechanisms including improving cerebrovascular health, lowering inflammation, reducing neurodegeneration, and improving brain metabolism and neuroplasticity.(46,47) Conversely, sedentary behavior is linked to increased risk of ADRD and cognitive decline.(48) There is some evidence that PA in humans may reduce the neurodegenerative aspects of dementia onset, with higher levels of PA associated with a lower abundance of neurofilament light chain (plasma marker of neurodegeneration), though the effect of PA on tau plaques and amyloid beta (Aβ) has had some conflicting results.(47,49)

Specific exercise prescriptions, which exercises provide what benefits, what is the optimal volume/ intensity/ duration, what factors distribute the exercise effect systemically, which exerkines cross the blood-brain barrier, and other questions remain regarding the relationship between exercise, age-related cognitive decline, MCI, and ADRD. Future work is needed to address these gaps in knowledge. As there have been both links and controversies between exercise intensity (i.e., higher = better) and multiple beneficial brain/cognitive adaptations, we chose to focus on the less well-studied HIIT paradigm.(50) In our current work we have demonstrated that late life enacted HIIT is sufficient to improve cognitive function, particularly in executive function tasks, in older adult female mice, with important translational potential for future investigations.

Aging is associated with chronic low-grade neuroinflammation characterized by persistent glial activation, increased proinflammatory cytokine signaling, and progressive disruption of neuronal homeostasis, particularly within hippocampal regions critical for cognition such as the CA1.(51–53) Recent studies suggest that neuroimmune aging trajectories differ by sex, with females demonstrating heightened inflammatory microglial responses during aging.(52,54) Despite this, most exercise-neuroimmunology studies remain heavily male-focused. Consistent with established aging phenotypes, sedentary aged females in our study exhibited elevated GFAP and IBA1 expression within the hippocampus. HIIT improved key features of glial morphology, including reduced astrocytic hypertrophy and increased microglial branching complexity, suggesting partial restoration of neuroimmune homeostasis even in the absence of significant changes in overall GFAP or IBA1 abundance. Given that glial morphology is increasingly recognized as an indicator of functional state, these findings suggest HIIT may preserve aspects of glial homeostasis in the aging female brain even in the absence of complete reversal of expression-based markers. In contrast to the modest structural glial changes, HIIT significantly reduced hippocampal IL-1β expression, identifying inflammatory signaling as a potentially early and highly exercise-responsive adaptation during aging. IL-1β is a central mediator of age-associated cognitive decline, synaptic dysfunction, and impaired hippocampal plasticity, with elevated signaling linked to deficits in learning and memory across both aging and neurodegenerative conditions.(54,55) Prior studies have shown that exercise can attenuate neuroinflammatory signaling during aging, although reported effects on glial activation markers remain inconsistent across exercise paradigms, durations, and sexes.(56–59) Our findings extend this by demonstrating that inflammatory cytokine signaling can be attenuated even when traditional glial markers remain elevated, suggesting that modulation of inflammatory pathways may precede broader cellular remodeling. These findings support the concept that HIIT preserves cognitive health during aging, at least in part, through modulation of hippocampal neuroimmune signaling in females.

### Cardiovascular Function

It is well-established that endurance training promotes beneficial cardiovascular adaptations in both human and animal models, across sexes and throughout the lifespan.(16,30,32) The gold standard of cardiovascular fitness is the VO_2_max test and endurance training increases this measure in most populations.(32) However, there are many other measures of cardiovascular fitness such as blood pressure, blood flow, heart pumping function, and ventricular wall thickness.

While there are numerous varieties of endurance exercise modalities used to promote cardiovascular fitness (i.e., swimming, running, walking, cycling, rowing), protocols using the modalities tend to gravitate towards a few different types: high volume/low intensity, moderate intensity continuous training (MICT), high intensity interval training, or sprint training (aka Wingate training).(16,60) There is some evidence that HIIT is equivalent, or even superior, to MICT for cardiovascular adaptations in humans with some variations depending on outcome markers, though MICT continues to be much more widely investigated.(16,61)

There is controversy over which type of training is better for blood pressure (BP) reduction, though the consensus is that endurance training overall improves BP versus sedentary behavior. For example, HIIT was shown to be superior to MICT in reduction of both systolic and diastolic blood pressure.(62) On the other hand, the complete opposite has also been reported for those *already with hypertension*, though HIIT improved overall cardiac fitness more.(63) In other studies of patients with hypertension, HIIT has been shown to improve overall cardiovascular fitness more than MICT (VO_2_max) while having at least equal effect on systolic and improved effect on diastolic BP.(64) In our current study, we saw an excellent improvement versus sedentary mice in both systolic and diastolic BP. Although the relative superiority of HIIT versus MICT for BP reduction remains unresolved, our data suggest that HIIT can produce meaningful antihypertensive benefits while also improving cardiovascular fitness-related outcomes.

As far as cardiac remodeling is concerned, there is also controversy over the effect of different endurance training types, intensities, and durations; and furthermore in some cases *sexual dimorphisms* result in differential adaptations.(16,65–67) In a study by Arbab-Zadeh and colleagues over a 12 month period of progressive endurance training in preparation to run a marathon, a group of sedentary individuals were measured for cardiac remodeling at baseline and then in 3-m intervals, with significant changes to ventricular (right and left) mass, and right diastolic volume, beginning at three months, though left ventricular end diastolic volume did not increase until after 6-9 months.(68) The authors noted that exercise intensity may be related to heart growth stimulus since the most intense exercise sessions were performed in months 7-9 of the study. In our study, we did not observe changes in left ventricular diastolic volume, which potentially could be due to the shorter duration of our protocol (14 weeks) and thus the stimulus may need to be longer to elicit volumetric alterations. However, we did observe a large effect size for increased ventricular wall thickness in the SED group, though this did not reach statistical significance.

In addition to functional cardiovascular adaptations, aging is associated with progressive extracellular matrix remodeling, fibroblast activation, and collagen accumulation that contribute to ventricular stiffening, impaired relaxation, and increased susceptibility to heart failure with preserved ejection fraction in older populations.(69) Both human and animal studies demonstrate age-associated increases in left ventricular fibrosis and hypertrophy, with female hearts showing pronounced collagen accumulation during aging.(70–72) Although endurance exercise is increasingly recognized as an effective strategy to attenuate myocardial fibrosis through mechanisms involving suppression of pro-fibrotic signaling pathways such as TGFβ1-Smad2/3, most preclinical studies have focused on males or moderate-intensity continuous training paradigms. In the present study, sedentary aged females demonstrated increased collagen deposition within the left ventricle, whereas HIIT attenuated this response, with exercised mice no longer differing from young controls.(70,73) These findings extend the current literature by demonstrating that HIIT is sufficient to blunt age-associated extracellular matrix remodeling specifically in the aging female heart. Previous studies report mixed effects of exercise on cardiomyocyte hypertrophy and fibroblast activation depending on exercise modality, duration, and disease context.(74,75) Here, αSMA expression and cardiomyocyte cross-sectional area trended higher in sedentary aged mice but were not evident in the exercised mice, suggesting partial suppression of maladaptive remodeling. This is important given that physiological exercise adaptations differ fundamentally from the pathological hypertrophy associated with aging and cardiovascular disease, where fibrosis and myofibroblast activation are major drivers of dysfunction. Interestingly, Cx43 expression remained unchanged despite evidence linking reduced Cx43 and enhanced fibrosis to aging-associated electrical remodeling and arrhythmogenic susceptibility.(76,77) This suggests that HIIT in aged females may preferentially target inflammatory and extracellular matrix remodeling pathways before overt electrophysiological remodeling becomes apparent.

### Caveats

The current study measures only a single age group, strain, and sex (female C57BL/6J, 24m old at endpoint), thus confirmation of these findings in similar age groups of males and other strains, as well as in older and younger groups, is required. Our previous work demonstrated that overall physical function is preserved by HIIT in adult (10m at study end) and older adult (24m) males, but neither cognitive nor cardiovascular function was measured in that study.(5) Earlier work with younger middle-aged males (17m old at study completion) did not show a robust increase in cognitive or physical function (other than in aerobic capacity), although the volume and intensity *was lower* than in the current study and we changed some of the cognitive function measuring parameters, so direct comparisons between these two studies are difficult.(14) In that study we hypothesized that since there was no evidence of cognitive decline over the period of the study in either the HIIT or SED group of middle-aged mice, the exercise stimulus was insufficient to improve cognition in that age-group and there was no deficit that could be prevented—in essence the 17m male mice were cognitively healthy. Notably, in both the current and previous work we found no change in the long-term memory version of the novel object recognition test, and we believe that the long-term test is not sensitive enough to detect short-term (3-4m) changes in cognitive ability in wild type non-pathological mice (i.e., normal aging), thus in the future we will use the “medium-term memory” NOR test where the second round is done ∼3 hours after the first instead of 24 hours and expect to see differences that we could not detect with the long-term memory test.

## Conclusions

HIIT exerted broad protective effects across multiple physiological systems in aged female mice, preserving or improving functional outcomes that typically decline with advancing age. Furthermore, we obtained evidence of positive effects on brain and heart health. Especially intriguing was the exercise-induced reduction of collagen deposition resulting in the restoration of heart fibrosis levels in the older females to similar levels as the young adult mice, while also vastly lower than that of the age-matched sedentary control. These findings position HIIT as a promising geroprotective intervention with the potential to preserve functional capacity, reduce frailty, and mitigate age-associated hallmarks of aging such as increased inflammation, fibrosis, and metabolic dysfunction. In the future we will need to extend these findings to different strains, other age groups of females and revisit males over the lifespan to measure cognitive and cardiovascular results.

Finally, we need further research to link specific mechanisms to the functional outcomes and to decode the vast network of interorgan signaling by exerkines and extracellular vesicles. Such studies may ultimately enable the development of novel therapeutic exercise mimetics that could reproduce select exercise benefits in those who cannot exercise either temporarily (in traction for broken hip) or permanently (quadriplegic).

## Supporting information

Online Only Supplmentary Section

## Acknowledgements

The authors wish to acknowledge Annika Bhardwaj and Robin Allgood for technical assistance. We also acknowledge the East Carolina Obesity and Diabetes Institute for their role in providing resources and equipment for this study (e.g., EchoMRI).

## Animal Use Statement

All animals were treated humanely under East Carolina University IACUC guidelines.

## Author Contributions

listed in order of contribution, **\*** = equal contribution The authors recognize the relative contributions as: **Conceptualization,** TGG, SS, LCB**; Methodology,** TGG, DT, SS, LCB, TT**; Validation,** TGG, SS, LCB, TT**; Formal Analysis,** TGG, DT, XS, AD, LT, LCB, AAO, AJ, SS**; Investigation** TGG, DT, PW, LT*, AAO*, AJ*, AD\***; Resources** TGG, SS, LCB, TT**; Writing – Original Draft** TGG, DT, AJ**; Writing – Review & Editing** TGG, DT, SS, LCB, TT, AD, XS*, LT*, PW*, AAO*, XS\***; Supervision** TGG, SS*, LCB\***; Project Administration** TGG, SS*, LCB\***; Funding Acquisition** TGG, SS, LCB, DT*, AAO*

