## Supplementary material for "High Intensity Interval Training in Aged Female Mice Preserves Physical, Cognitive, and Cardiovascular Function": Online Only Supplmentary Section

**Table of Contents**

| Section | Page # |
|---|---|
| Supplemental Methods Section | S2 |
| Table S1 SOL IHC Antibodies | S5 |
| Supplemental Results Section | S6 |
| Figure S1 Running Speed | S10 |
| Figure S2 Body Composition | S11 |
| Figure S3 Whole Body Calorimetry | S12 |
| Supplemental References | S13 |
| Appendix A: Frailty Index Scoring Sheet | S15 |
| Dataset S1 CFAB Details | Excel File |
| Dataset S2 CAB Details | Excel File |
| Dataset S3 Cardiovascular Details | Excel File |
| Dataset S4 Body Composition Details | Excel File |

**Supplemental Methods**

*Functional Testing:*

*CAB (Cognitive Assessment Battery):* We converted the CAB determinant score into a composite score by determining which quartile an individual mouse was in for each of our included measures: open field total distance and center time, NOR discrimination ratio and total exploration time, puzzle box total time (challenge 1 and 2) and total time challenge 3, and y-maze total alternations and spontaneous alternation ratio, then summed these to get a total score, with higher score equal better performance. Note that in the puzzle box tests quartiles were in reverse order since lower numbers were better (i.e., quartile 1 was the best, so it was converted to 4 to match the other tests and so forth), and in the other three tests higher numbers were better (i.e., quartile 4 is best).

*Frailty:* We used a combination of previous methods to determine frailty.[1-3] The five determinants for the pre- and post-training measurements were: 1) volitional activity rate (voluntary wheel running in km per day, averaged over the time spent in the Promethion metabolic cages that housed the wheels.), 2) overall motor function (measured with rotarod as latency to fall in seconds), 3) forelimb grip strength (measured by grip meter in milliNewtons per gram of body mass), 4) whole body strength/endurance (inverted cling test, measured as latency to fall in seconds), and 5) aerobic capacity and endurance (measured by the treadmill max speed test in seconds performed). We then also analyzed only the post-intervention groups using a modified version of the Frailty Index (FI) as a 6$^{th}$ determinant as we recently published.[3-5] We thus had two scores, one with 5 determinants measured pre- and post-intervention and a second score where we added the FI component as a sixth determinant. To assess frailty, for each determinant we calculated a cut-off point that was 1.5 standard deviations from the mean of the entire group (both HIIT and SED) at baseline (other than the FI which was only measured post-training and thus was indexed against the mean frailty score at that time point). Any mouse that was beyond the cut-off point received a score of 1, otherwise they scored zero for that determinant. We then summed the total score for all determinants and mice scoring 3 or more were considered frail, 2 as pre-frail, and 1 or 0 as non-frail.

*Blood pressure measurement:* We used noninvasive tail-cuff plethysmography according to the manufacturer's protocol (CODA Kent Scientific). Prior to baseline blood pressure measurements, mice acclimatized to the procedure for three consecutive days. All experiments were performed in a designated quiet area at the same time each day. Mice entered appropriately sized restraint tubes, and nose cones adjusted to minimize movement. Heating platforms maintained a body temperature of 32°-35°C before and throughout recordings. We inspected the occlusion and volume pressure recording (VPR) cuffs for leaks prior to each session. The occlusion cuff was positioned at the base of the tail with the VPR sensor placed directly behind it. The occlusion cuff was inflated and deflated over 20 s while the VPR sensor detected changes in the tail volume during blood return. The minimum tail volume was set to 15 µL. Mice underwent 5 acclimation cycles followed by 25 measurement cycles with an 8 s interval between cycles. We used at least 5 accepted values from the measurement cycles for analysis. We recorded blood pressure for three consecutive days each week and averaged the values to obtain one measurement for each mouse.

*Echocardiography imaging:* Echocardiography imaging is a non-invasive procedure that allows assessment of both systolic and diastolic function. We performed echocardiography twice for each animal; before training and after training, in both the HIIT and SED groups, using a digital ultrasonic imaging system (Vevo 3100, Fujifilm VisualSonics, USA). Briefly, mice were initially anesthetized by placing them in a flow-through system containing 2-3.5% isoflurane in 100% oxygen. Following loss of consciousness, we placed the mice on a modified mask assembly that allowed continuous delivery of 1-2.5% isoflurane in oxygen. The mice breathed spontaneously, and the operator monitored the depth of anesthesia throughout the continuous recording of heart rate. It was crucial to maintain heart rate above 400 beats per minute, as the accuracy of the images obtained depends on a near physiological heart rate. Mice acclimatized for ~5min on an isothermal pad to maintain body temperature, and then hair removed where needed before recording images, as described previously and according to the Guidelines for Measuring Cardiac Physiology in Mice.[6-8]

*Body Composition:* We measured body mass weekly and body composition (with EchoMRI) pre- and post-intervention. We have previously detailed EchoMRI, which we use to report lean mass and fat percentage changes.[9,10]

*Whole Body Metabolism:* We utilized Promethion Metabolic Chambers (Sable Systems) by following the manufacturer's instructions at a 12-hour/12-hour light/dark cycle held at a constant 23°C for 6 or 7 days. Both the running wheels and the laser x/y/z plane activity grids were active to measure voluntary wheel running and activity rate, respectively and adjusted to meters per day (m/day). We report whole body calorimetry (vO2, oxygen consumption, vCO2, carbon dioxide production, RER, respiratory exchange ratio, and kilocalories per hour expended, Kcal/hr) as the overall mean and the maximum number recorded. Food and water consumption, and body mass, were recorded throughout the length of the procedure (data not shown).

<u>*Ex Vivo Outcome Measures:*</u>

*Immunohistochemistry:* We collected brain and heart tissues from anesthetized mice and post-fixed them in 4% paraformaldehyde in 0.1 M phosphate-buffered saline (PBS; pH 7.4), followed by cryoprotection in 30% sucrose in PBS overnight. We sectioned brains coronally at 25 µm using an HM 430 sliding microtome. After embedding in paraffin, we sagittally sectioned the hearts at 5 µm onto glass slides. Paraffin sections were deparaffinized and subjected to heat-induced antigen retrieval. Immunofluorescence staining was performed, as previously described.[11] Briefly, free-floating brain sections and heart sections were washed in 1× PBS and incubated for 1 h at room temperature in blocking solution containing 2% normal donkey serum, 0.05% Tween-20, 50 mM glycine, 0.1% Triton X-100, 0.1% bovine serum albumin, and 1x PBS. Sections were then incubated overnight at 4 °C in a humidified chamber with the following primary antibodies: connexin 43 (Cx43; Sigma-Aldrich C6219, 1:500), α-smooth muscle actin (αSMA; Novus NB300-978, 1:500), glial fibrillary acidic protein (GFAP; Invitrogen 14-9892-2, 1:500), AIF1/IBA1 (ABclonal A19776, 1:1000), and interleukin-1β (IL-1β; Invitrogen P420B, 1:500). Following washes in PBS containing 0.5% Tween-20, sections incubated for 2 h at room temperature with the appropriate Alexa Fluor conjugated secondary antibodies (Invitrogen A32794, A32849, A32816, A32795; all 1:1000). We then mounted sections using VECTASHIELD antifade mounting medium with DAPI (Vector Laboratories, H-1800). We captured a minimum of five images per section using a fluorescence microscope, and quantified mean fluorescence intensity using ImageJ software (NIH). We selected the representative images from samples whose quantified values were closest to the group mean to best reflect the average response.

*Cardiomyocyte Cross-Sectional Area:* We labeled, the cardiomyocyte cell membranes with wheat germ agglutinin (WGA) conjugated to Alexa Fluor 488 (Invitrogen W11261; 1 mg/mL stock, 1:200 dilution) for 10 min as previously described.[12,13] We mounted sections with VECTASHIELD antifade mounting medium with DAPI. Images were acquired using a fluorescence microscope

(Keyence/Echo Revolve). A blinded investigator quantified cardiomyocyte cross-sectional area using ImageJ.

*Measurement of Collagen:* To determine the extent of collagen deposition as a measure of cardiac remodeling, we stained heart tissue sections with 0.1% Picrosirius Red to visualize fibrillar collagen as previously described.[8] Sections were imaged under standard brightfield microscopy by an investigator blinded to the experimental groups. The collagen-positive staining was quantified using identical thresholding parameters across all groups.

*Microglial Morphological Analysis*: We quantified microglial reactivity through AIF1/IBA1-labeled images acquired at 63x magnification using a confocal microscope. We analyzed microglial morphology using the AnalyzeSkeleton (2D/3D) plugin in Fiji/ImageJ, as previously described, images were converted to 8-bit grayscale, and brightness and contrast were adjusted using the maximum slider to visualize microglial processes.[11] Contrast was further enhanced using the Unsharp Mask tool (pixel radius: 3; mask weight: 0.6), followed by despeckling to reduce noise. Images were converted to binary, skeletonized, and analyzed to quantify average length and number of branches.

*Astrocyte Ramification Analysis*: We quantified astrocyte complexity using the Shoenen ramification index, defined as the ratio of the maximum number of Sholl intersections to the number of primary branches. Images were imported into Fiji and analyzed using the Simple Neurite Tracer (SNT) plugin. Astrocytes were reconstructed according to the Fiji SNT walkthrough protocol (https://imagej.net/plugins/snt/walkthroughs), with the starting point defined as the center of DAPI staining and all primary processes traced outward. We performed Sholl analysis using the Fiji SNT Sholl protocol (https://imagej.net/plugins/snt/sholl) at a radius step size of 4 µm.

*Soleus (SOL) Muscle Morphology:* As previously described, we used immunofluorescence to test for fiber type distribution (myosin heavy chain 1, MHC1, MHC2a, MHC2x, and hybrid fibers) and cross-sectional area by fiber type in a randomly selected subset of mice (n=6 for SED and n=4 for HIIT).[14,15] See expanded methods section in the online supplement for further details, including antibodies and concentrations used. Briefly: We cut frozen sections of SOL at 7 µm thickness on a cryostat, mounting 3 SOL sections per slide. The sections were airdried for 1 hour, then rehydrated in phosphate buffered saline (PBS), each section circled with a hydrophobic PAP pen, then immersed in MoM (mouse on mouse blocker, Vector Labs), primary antibodies for 90 minutes at RT (room temperature), then secondary antibodies for 60 minutes at RT, followed by post-fixing in methanol (see **Table S1** for antibodies). We washed slides with PBS washes between steps. We mounted coverslips using Vectashield antifade mounting medium (Vector Labs) and took images on an EVOS fluorescent microscope at 200x total magnification. Data was Blinded investigators quantified the data using ImageJ.

| Primary Antibody | Target | Supplier | Concentration | Secondary Antibody | Concentration |
|---|---|---|---|---|---|
| BA.D5 IgG2b mouse Concentrate[1] | **MHC1** | DHSB Iowa | 1:100 | Gt anti-Ms IgG2b, Alexa Fluor 647 | 1:500 |
| SC.71 IgG1, mouse supernatant[2] | **MHC2a** | DHSB Iowa | 1:4 | Gt anti-Ms IgG1, Alexa Fluor 488 | 1:250 |
| BF.F3 IgM, mouse supernatant[3] | **MHC2b** | DHSB Iowa | 1:4 | Gt anti-Ms IgM, Alexa Fluor 555 | 1:250 |
| Anit-laminin IgG Rabbit Polyclonal | **laminin** | Sigma #L9393 | 1:200 | *Gt anti-Rb IgG, Alexa Fluor 350* | 1:250 |

**Table S1: Soleus IHC Antibodies. KEY:** DHSB Iowa = Developmental Studies Hybridoma Bank

**NOTES:**
[1] The BA.D5 IgG2b developed by S. Schiaffino of the Universita degli Studi di Padova was obtained from the Developmental Studies Hybridoma Bank, created by the NICHD of the NIH, and maintained at The University of Iowa, Department of Biology, Iowa City, IA 52242.

[2]The SC.71 IgG1 mouse concentrate developed by S. Schiaffino of the Universita degli Studi di Padova was obtained from the Developmental Studies Hybridoma Bank, created by the NICHD of the NIH, and maintained at The University of Iowa, Department of Biology, Iowa City, IA 52242.

[3] The BF.F3 IgM developed by S. Schiaffino of the Universita degli Studi di Padova was obtained from the Developmental Studies Hybridoma Bank, created by the NICHD of the NIH, and maintained at The University of Iowa, Department of Biology, Iowa City, IA 52242.

**Supplemental Results**

<u>*In vivo Outcome Measures:*</u>

*Physical Function:* We measured physical function using the well-validated CFAB tests of rotarod, VWR, grip meter, inverted cling, and max speed test (treadmill), as well as x/y/z plane laser grid activity monitoring *(see* **Figure 2 Physical Function**). There were many changes within groups from pre- to post-training paired t-tests, but fewer between groups (independent t-tests). A comprehensive breakdown of the statistical analysis is below with specifics in **Supplemental Dataset S1. Table 1** in the main paper lists the more pertinent outcomes.

- Rotarod (overall motor function):
  - Within groups: rotarod time (SED: NC, no statistical change, numeric difference -14.1 ± %, p=0.373, Hedges' g 0.30; HIIT: +39.1% ± 14.9, *trend* p=0.078, Hedges' g 0.65).
  - Between groups: rotarod time (PRE: NC, p=0.528; Hedges' g 0.283, POST: NC, p=0.222; Hedges' g 0.61); rotarod percent change (NC, p=0.251, Hedges' g 0.59)
- VWR (volitional exercise):
  - Within groups: rotarod time (SED: -41% ± 25, p=0.004, Hedges' g 1.34; HIIT: -42% ± 23, *trend* p=0.002, Hedges' g 1.50).
  - Between groups: VWR meters per day (PRE: NC, p=0.528; Hedges' g 0.42; POST: *trend* p=0.097; Hedges' g 0.84); VWR percent change (NC, p=0.774, Hedges' g 0.148)
- Grip Meter (fore-limb strength):
  - Within groups: grip time (SED: -16% ± 4, p=0.007, Hedges' g 1.18; HIIT: NC, numeric +6% ± 12, p=0.838, Hedges' g 0.067); grip_mass (grip adjusted by multiplying with grams body mass) (SED: -35% ± 8, p=0.002, Hedges' g 1.56; HIIT: -NC. Numeric -14% ± 9, p=0.184, Hedges' g 0.65)
  - Between groups: grip time (PRE: NC, p=0.905; Hedges' g 0.424, POST: p=0.043; Hedges' g 1.05); grip time percent change (NC, p=0.126; Hedges' g 0.799); grip_mass (grip adjusted by body mass) (PRE: NC, p=0.822; Hedges' g 0.098, POST: *trend* p=0.053; Hedges' g 1.00); grip_mass percent change (PRE: trend *p=0.104*; Hedges' g 0.424, POST: p=0.043; Hedges' g 1.05)
- Inverted Cling (four limb strength/endurance):
  - Within groups: cling time (SED: -67% ± 6, p=0.004, Hedges' g 1.35; HIIT: -37% ± 10, p=0.025, Hedges' g 0.89); cling * mass (work) (SED: -64% ± 6, p=0.002, Hedges' g 0.49; HIIT: -14% ± 9, *trend* p=0.052, Hedges' g 0.46)
  - Between groups: cling time (PRE: NC, p=0.356; Hedges' g 0.16, POST: *trend* p=0.058; Hedges' g 0.1.00), cling * mass (PRE: NC, p=0.766, Hedges' g 0.283, POST: p=0.043, Hedges' g 1.05), cling time percent change (NC, p=0.126, Hedges' g 0.80); grip * mass percent change (SED -35%, HIIT -14%, *trend* p=0.104, Hedges' g 0.82)
- Treadmill (aerobic capacity, endurance):
  - Within groups: treadmill time (SED: NC, p=0.381, Hedges' g 0.30; HIIT: +71% ± 17, p=0.002, Hedges' g 1.56).
  - Between groups: treadmill time (PRE: p=0.528; Hedges' g 0.283, POST: p=0.028; Hedges' g 1.16); treadmill percent change (SED +6%, HIIT +71%, p=0.005, Hedges' g = 1.75)
- Activity Monitoring (activity rate):
  - Within groups: total meters traveled per day (SED: NC, p=0.109, Hedges' g 0.58; HIIT: NC, p=0.881, Hedges' g 0.65).

  - Between groups: total meters traveled per day (PRE: NC, p=0.393; Hedges' g 0.370, POST: p=0.739; Hedges' g 0.57)

*Cognitive Function:* We measured cognition using open field, NOR, y-maze, and puzzle-box *(see* **Figure 4** and **Supplemental Dataset S2 Cognitive Function** for more details). Differences in means within-groups were measured with paired t-test, and between groups were measured with Student's independent samples t-test.

- Open Field (exploratory behavior and anxiety):
  - Within groups multiple parameters changed pre- to post-intervention in both groups including: total distance traveled (SED: +75.0% ± 41.5, p=0.005, Hedges' g 0.52; HIIT: +89.3% ± 28, p<0.001, Hedges' g 1.36), average speed (SED: +74.6% ± 50.0, *trend* p=0.067, Hedges' g 0.53; HIIT: +90.1% ± 44.6, p<0.001, Hedges' g 1.36), maximum speed (SED: +23.6% ± 9.1, p=0.012, Hedges' g 0.53; HIIT: +50.6% ± 20.6, p=0.026, Hedges' g 0.79), time immobile defined as not moving for ≥ 10 seconds (percent change of overall mean pre- to post: SED: -88.6%, *trend* p=0.090, Hedges' g 0.56; HIIT: -89.9%, p=0.004, Hedges' g 1.21), and number of total immobile episodes (percent change of overall mean pre- to post: SED: -75%, *trend* p=0.052, Hedges' g 0.55; HIIT: -86.1%, p=0.001, Hedges' g 1.39). Anxiety was significantly reduced in HIIT, but not SED, reflected by increased time in center of field (SED: +97.8% ± 9.1, p=0.012, Hedges' g 0.49; HIIT: +50.6% ± 20.6, p=0.026, Hedges' g 1.03).
  - Between groups: There were no significant differences either before or after training.
- NOR (long-term memory):
  - Within groups: Multiple parameters changed pre- to post-intervention, including: total distance traveled (SED: +88.3% ± 34.1, p=0.005, Hedges' g 1.25; HIIT: +93.3% ± 41.3, p=0.006, Hedges' g 1.22), average speed (SED: +90.0% ± 34.5, p=0.005, Hedges' g 1.26; HIIT: +92.3% ± 40.2, p=0.006, Hedges' g 1.22), maximum speed (SED: +26.2% ± 3.9, p<0.001, Hedges' g 2.10; HIIT: +28.7% ± 6.2, p=0.006), time immobile (SED: -49.3% ± 8.8, p=0.016, Hedges' g 0.98; HIIT: -59% ± 14.0, p=0.007, Hedges' g 1.17), number of total immobile episodes (percent change of overall mean pre- to post: SED: +50.8%, p=0.004, Hedges' g 1.35; HIIT: -52.8%, p=0.014, Hedges' g 0.83). There was evidence for trends in changes from pre- to post-training to the investigation time of the novel object (NO) in both groups but there were no significant changes in familiar object (FO) investigation time: NO (SED: +50.8% ± 29.7, *trend p=0.053*, Hedges' g 0.73; HIIT: +300.4% ± 190.0, *trend p=0.096*, Hedges' g 0.53), FO (SED: +191% ± 119.2, p=0.11, Hedges' g 0.23; HIIT: +221.2% ± 149.9, p=0.822, Hedges' g 0.23); with obvious large individual variability. However, the major indicator of long-term memory, the discrimination index (measures the time spent investigating the familiar object versus time spent investigating the novel object) was not different between or within groups, either before or after, the intervention period.
  - Between groups: There were no significant differences either in pre- or post-intervention, in part due to large individual variability.

- Puzzle Box (executive function and memory):
  - Within groups: multiple parameters changed pre- to post-intervention including: PB1 (Puzzle Box parameter 1, the time to entry of escape trial on day1; SED: -45.2% ± 13.1, p=0.01, Hedges' g 1.1; HIIT: -68.3% ± 22.1, p=0.001, Hedges' g 1.6), PB2 (Puzzle Box parameter 2, the time to entry of escape trial on day 2; SED:

no change, *NS* p=0.411 Hedges' g 0.27; HIIT: -93.1% ± 1.9, p=0.021, Hedges' g 0.93), PB1_2total (sum of Puzzle Box parameters 1 and 2; SED: -45.0% ± 12.1, p=0.015, Hedges' g 1.01; HIIT: -75.6% ± 4.3, p=0.004, Hedges' g 1.33), PB3.1 (time to entry of escape trial on day 3, measured from trial start; SED: -46.5% ± 16.1, p=0.043, Hedges' g 0.78; HIIT: -82.1% ± 5.5, p=0.005, Hedges' g 1.26), PB3.2 (time to clearing blockage to escape box on day 3, measured from trial start; SED: no change, p=0.144, Hedges' g 0.52; HIIT: -45.2% ± 12.5, p=0.011, Hedges' g 1.08), PB3_total (sum of PB3.1 and PB3.2 on day 3; SED: -28.5% ± 17.8, *trend* p=0.065, Hedges' g 0.69; HIIT: -60.0% ± 9.8, p=0.005, Hedges' g 1.25), and PBtotal (sum of PB1, PB2, PB3.1 and PB3.2; SED: -40.4% ± 13.3, p=0.021, Hedges' g 0.93; HIIT: -72.8% ± 3.0, p=0.002, Hedges' g 1.50).
    - Between groups: There were no significant differences in pre-intervention. However, there were significant differences post-training found in PB1_2 percent change (p=0.032), PB3.1 percent change (*trend*, p=0.055), and in the overall major indicator of executive function PBtotal percent change (p=0.032), all of which demonstrated a greater effect in the HIIT group.

- Y-maze (spatial memory and exploratory behavior):
    - Within groups: Multiple parameters changed pre- to post-intervention including: total distance traveled (SED: no change, p=0.209, Hedges' g 0.44; HIIT: +45.6% ± 9.2, p=0.002, Hedges' g 1.94), average speed (SED: no change, p=0.217, Hedges' g 0.43; HIIT: +45.6% ± 9.3, p=0.002, Hedges' g 1.90), maximum speed (SED: +13.6% ± 5.7, *trend* p=0.064, Hedges' g 0.69; HIIT: +35.5% ± 4.3, p<0.001, Hedges' g 4.23), time immobile (SED: no change, p=0.179, Hedges' g 0.66; HIIT: -61.0% ± 7.5, p=0.011, Hedges' g 1.23), number of immobile episodes (SED: no change, p=0.487, Hedges' g 0.30; HIIT: -86.1%, p=0.003, Hedges' g 1.56), total arm entries (SED: +44.9% ± 14.5, p<0.001, Hedges' g 1.11; HIIT: +58.1% ± 5.4, p<0.001, Hedges' g 3.64 ), and number of spontaneous alternations (SA) defined as traveling subsequently into 3 different arms (e.g., AàBàC) without backtracking as a measure of spatial/short term memory (SED: +65.9% ± 2.8, *trend* p=0.077, Hedges' g 0.71; HIIT: +56.8% ± 2.8, p=0.001, Hedges' g 1.63 ). However, the major y-maze indicator percent of SA in relation to total arm entries did not change (Hedges' g: SED 0.33, HIIT 0.49), as both total number of entries and number of SA increased proportionally.
    - Between groups: There were no significant differences in pre-intervention, other than related to average speed while traveling the maze (2.68 m/min in SED vs. 2.51 m/min in HIIT, p=0.080, Hedges' g: 0.89). There were no significant differences post-intervention other than percent change in maximum speed (+35.5% in HIIT versus +13.6% in SED, p=0.009, Hedges' g: 1.45) and percent change of immobile time (-61% in HIIT versus -30% in SED, *trend*, p=0.098, Hedges' g: 0.862), both demonstrating a stronger effect in HIIT.

*Functional Composite Scores:*

- <u>CFAB:</u> Comprehensive functional assessment battery composite scoring system for physical function determined that HIIT preserved and SED lost functional capacity. See **Figure 4** (main paper), **Table 3** (main paper), and **Supplemental Dataset** S1 for more information.

    - Within groups: CFAB (SED: Pre – Post = 4.52 ± 0.75, p<0.001, Hedges' g 0.30; HIIT: NC, numeric change Pre – Post = 0.77 ± 1.06, p=0.490, Hedges' g 0.41).

  - Between groups: no significant differences in CFAB (CFAB1 PRE: $p=0.528$; Hedges' g 0.283, CFAB2 POST: $p=0.222$; Hedges' g 0.61); but the changes in CFAB pre- to post were different: ΔCFAB ($p=0.021$; Hedges' g 1.23)

- CAB: Cognitive assessment battery composite scoring system determined cognitive function improved with HIIT but not in SED. See **Supplemental Dataset S2.**
  - Within Groups CAB (SED: no change, $p=0.921$, Hedges' g 0.03; HIIT: +25%, $p=0.022$, Hedges' g 0.92).

  - Between groups: no significant differences in CAB raw scores (CAB1 PRE: $p=0.391$; Hedges' g 0.42, CAB2 POST: $p=.240$; Hedges' g 0.61); but the changes to CAB from pre- to post were different: ΔCAB ($p=0.039$; Hedges' g 1.08), CAB_percentchange (SED +1.99% versus HIIT +34.08%, $p=0.067$; Hedges' g 0.98).

*Body Composition:* See **Figure S3 Body Composition** and **Supplemental Dataset S4 Body Composition** for further details. We tracked body mass monthly during aging up to training and then weekly once training commenced. We measured fat percentage and total lean mass totals using EchoMRI, pre- and post-training.

- Within groups:
  - Body mass increased in both groups from pre- to post-training (SED: +9%, *trend* $p=0.063$, Hedges' g 0.69; HIIT: +12.8%, $p=0.009$, Hedges' g 1.12).
  - Fat mass (SED: NC, numerical +2.69 g ±1.57, $p=0.110$, Hedges' g 0.58; HIIT: +3.26 g ± 0.88, $p<0.001$, Hedges' g 1.16). However, SED fat mass post-training had a minor violation of normality (Shapiro-Wilk sig.=0.032), and when the pre/post means were analyzed with the Wilcoxon Signed Rank test, fat mass was significantly increased (sig. = 0.017).
  - lean mass (SED: +0.95 g ± 0.23, $p=0.005$, Hedges' g 1.28; HIIT: +0.65 g ± 0.11, $p<0.001$, Hedges' g 1.87)
  - Fat percentage (SED: NC, numerical +5.1% ± 3.5, $p=0.183$, Hedges' g 1.11; HIIT: +6.6% ± 1.7, $p=0.006$, Hedges' g 1.23).

- Between Groups: There were no significant changes between groups either pre- or post-training in body mass, lean mass, fat mass, or fat percentage.

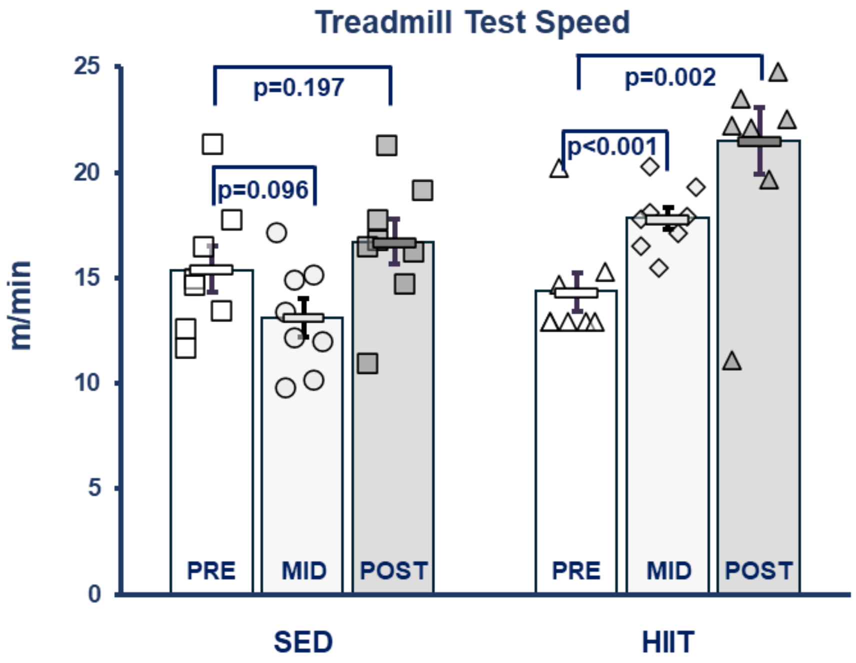

**Figure S1 Treadmill Speed.** A 2x3 (2 groups, SED and HIIT x three times, PRE, MID, and Post) mixed-model ANOVA showed with-in subject main effects of time ($F=17.07$, $p<0.001$, $\eta^2=0.549$) and the group*time interaction ($F=9.15$, $p<0.001$, $\eta^2=0.395$), and differences in means between subjects ($F=5.252$, $p=0.038$, $\eta^2=0.273$). Over the course of the training period the maximum treadmill speed of the SED mice did not significantly alter (paired t-test: pre->mid, p=0.096, Hedge's g= 0.604; pre->post, p=0.222, g=0.448), though there was a trend of a decline at midpoint. The HIIT mice had significant increases in maximum treadmill speed at both the midpoint (+18% at 7 weeks, p<0.001, g=1.96) and post-training (+26% at 14 weeks p=0.002, g=1.56) compared to the baseline pre-training speed. At baseline Sed and HIIT had no difference between groups (p=0.470, independent samples t-test), but HITT had faster times at both the midpoint (+36%, p<0.001, g=2.11) and the post-training tests (+29%, p=0.028, g=1.56), indicating positive aerobic capacity adaptations from training. KEY: SED=sedentary control group, HIIT=high intensity interval training group, PRE= before intervention period, MID=middle time point, POST=after intervention, squares/circles=SED, triangles/diamonds=HIIT, each symbol=speed of one mouse. Units are meters per minutes (m/min). The p-values in graph are from paired t-tests with pre. Rectangles = means per group with standard error bars.

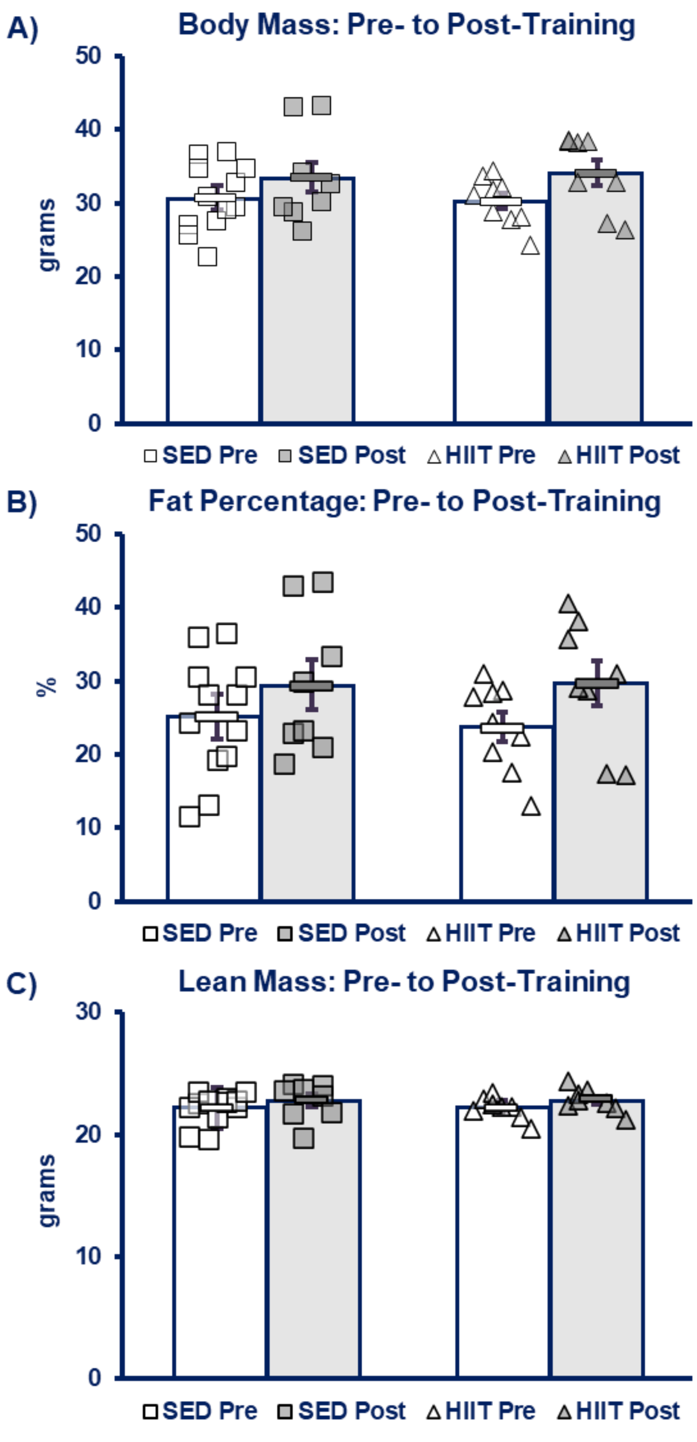

**Figure S2 Body Composition. T**here were no differences between groups (independent samples t-test). **A) Body Mass. B) Fat Percentage. C) Lean Mass. KEY:** SED=sedentary control group, HIIT=high intensity interval training group, PRE=before intervention period, POST=after intervention, squares=SED, triangles=HIIT, open symbols=pre, shaded symbols=post, each symbol=one mouse. The p-values in graph are from paired t-tests. Rectangles = means per group with standard error bars. Percentages in bar graphs = percent change from per- to post intervention.

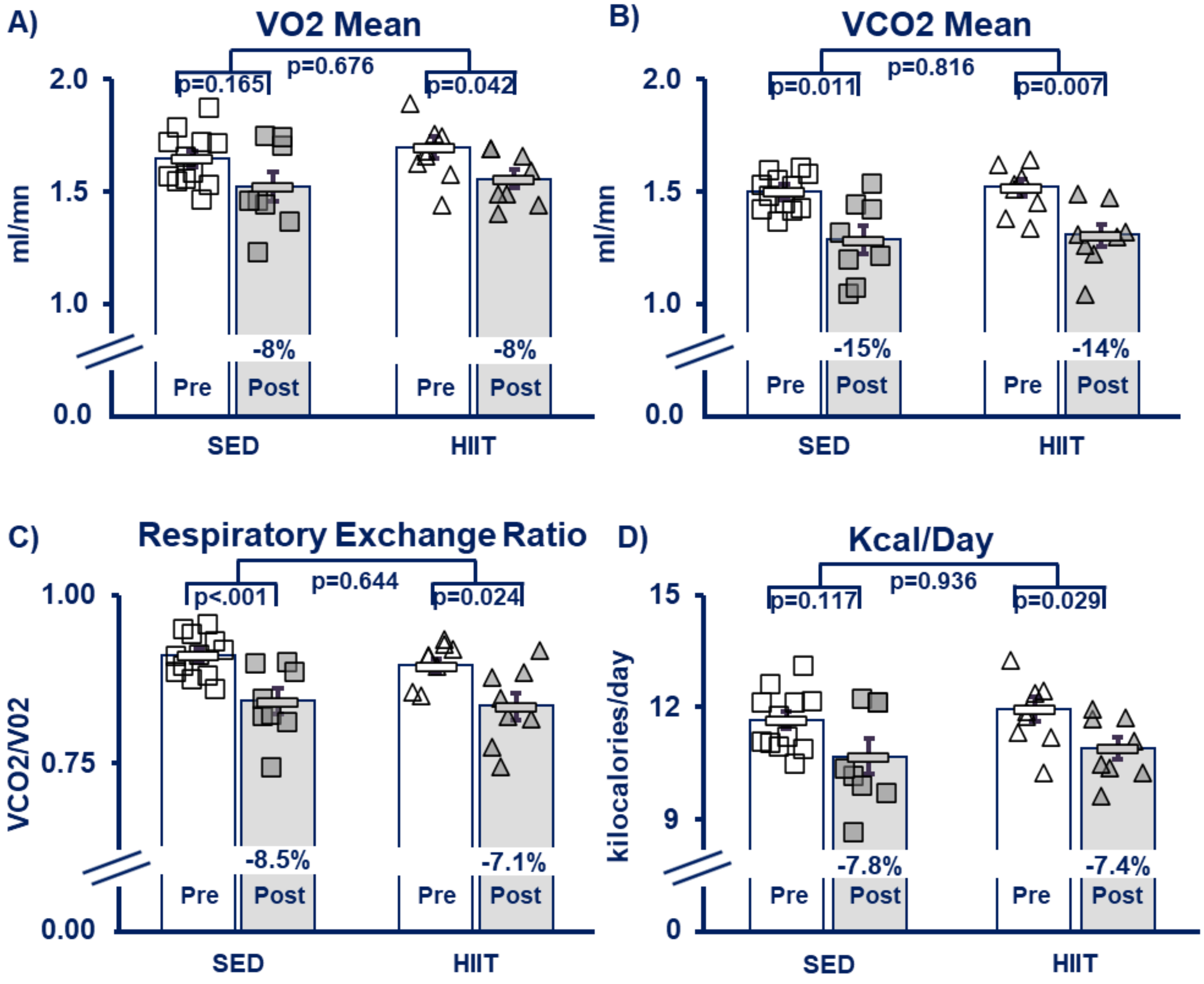

**Figure S3 Metabolic Chamber Outcomes. A) VO2.** Average oxygen consumption. **B) VCO2.** Average Carbon dioxide production. **C) Respiratory Exchange Ratio. D) Average kilocalories burned per day. KEY:** SED=sedentary control group, HIIT=high intensity interval training group, PRE= before intervention period, POST=after intervention, squares/circles=SED, triangles/diamonds=HIIT, each symbol=one mouse. The p-values in graph are from paired t-tests with pre. Rectangles = means per group with standard error bars.

### Appendix A: Sample Frailty Index Form

#### Frailty Index Criteria Sheet

| Group | Animal ID & FI Total |
|---|---|
| Date | |
| Alopecia | |
| Loss of fur color | |
| Dermatitis | |
| Loss of whiskers | |
| Coat condition | |
| Tumours | |
| Distended abdomen | |
| Kyphosis | |
| Tail stiffening | |
| Gait disorders | |
| Tremor | |
| Body condition score | |
| Vestibular disturbance | |
| Hearing loss | |
| Cataracts | |
| Corneal opacity | |
| Eye discharge/swelling | |
| Microphthalmia | |
| Vision loss | |
| Menace reflex | |
| Nasal discharge | |
| Malocclusions | |
| Rectal prolapse | |
| Vaginal/uterine/penile prolapse | |
| Diarrhea | |
| Breathing rate/depth | |
| Mouse Grimace Scale | |
| Piloerection | |
